# Modeling patient variants of *Cnot1* and *Cdc42bpb* results in distinct forms of congenital diaphragmatic hernia in mice

**DOI:** 10.64898/2026.02.24.707527

**Authors:** Eric L. Bogenschutz, Cynthia Carpenter, Ameleen Wong, Kristina Palmer, Aalekh Mehta, Yannick Ledermann, Caleb Heffner, Kathy J. Snow, Carol Bult, Yufeng Shen, Patricia K. Donahoe, Samuel P. Rowbotham, Frances A. High, Wendy K. Chung, Stephen A. Murray

## Abstract

Congenital diaphragmatic hernia (CDH) is a severe congenital anomaly characterized by impairment of both diaphragm and lung development *in utero*. CDH presents as a spectrum of forms and severities, with diaphragm defects arising in the dorsal/posterior region typically correlating with more severe pulmonary disease and higher risk of mortality than those appearing in ventral/anterior regions. The genetic etiology underlying CDH is complex, with many genes implicated showing variable expressivity and incomplete penetrance in both human patients and mouse models. Here we present *in vivo* validation of two genes previously unassociated with CDH: the CDC42-interacting kinase *CDC42BPB*; and *CNOT1*, a scaffolding protein of the CCR4-NOT protein complex, critical for mRNA regulation through modifications such as deadenylation. Each gene was found to have a damaging, *de novo* missense variant in a recent large-scale CDH patient sequencing screen. Loss of *Cdc42bpb* leads to ventral diaphragmatic hernias, heart septal defects and minor lung epithelial differentiation defects in mouse embryos. Installation of the orthologous patient-specific missense variant through CRISPR/Cas9 editing leads to less severe ventral diaphragm defects. Mouse embryos with either one or two copies of the orthologous *Cnot1* variant, c.1867C>T (p.R623W), develop dorsal diaphragmatic hernias with low (<50%) penetrance, and mutants showed alterations in mRNA isoform expression consistent with the molecular role of *Cnot1* in RNA splicing. These results underscore the power of *in vivo* functional modeling to validate genes and patient-specific variants uncovered by patient sequencing, reveal two previously unrecognized genetic causes of CDH, and highlight the heterogeneity of different patient anatomic presentations.

## Introduction

Congenital diaphragmatic hernia (CDH) is a relatively common and deadly congenital anomaly impacting multiple organs of the respiratory system. CDH is characterized by a weakness or partial absence of the diaphragm muscle, allowing abdominal contents to enter the thoracic cavity. It is associated with abnormal lung development and function due to both lung-intrinsic and -extrinsic (mechanical) factors (1, 2). CDH arises in approximately 1 in 2,500 live births and results in mortality in an estimated 30-50% of cases (1, 2). Mortality is due primarily to hypoplastic lung development with incomplete formation of alveoli and respiratory distress at birth (2, 3). Both human and animal studies have demonstrated CDH is caused by abnormal development of the embryonic diaphragm and lung, and that *de novo* genetic alterations account for a substantial portion of CDH cases (1, 4, 5).

Exome and genome sequencing efforts have revealed many gene variants associated with human CDH. These variants range in size, inheritance, and recurrence across patients, and include single nucleotide variants (SNVs), small insertion and deletions (indels) and large structural variants (SVs) (1, 6). However, only approximately 30% of cases will have a genetic diagnosis from genomic sequencing (4, 5). Although a number of causative genes for CDH have been identified and validated through rigorous statistical and/or functional assays, these ongoing discovery efforts continue to identify novel genes and variants, frequently in genes with limited functional annotation. Functional validation to corroborate genotype-phenotype relationships and to gain a better understanding of their biological function in CDH and normal diaphragm development is crucial for better diagnosis and clinical intervention in CDH.

The diaphragm is a sheet-like skeletal muscle that is unique to mammals and essential for survival due to its role in respiration (7). The diaphragm develops from two transient embryonic structures: the somites, located lateral to the neural tube that give rise to all skeletal muscle, and the pleuroperitoneal folds (PPFs), located on the dorsal anterior edge of the developing liver, that give rise to the connective tissue of the diaphragm (8). Muscle progenitors migrate from the somites into the PPFs around mouse embryonic day (E)11.5 (9), as the PPFs start to migrate dorsally and ventrally, spreading across the liver and the mesenchymal septum transversum (9). As the PPFs migrate, muscle progenitors differentiate into muscle fibers and neuronal projections from the phrenic nerve migrate and branch to form neuromuscular junctions (9, 10). In mice, the diaphragm is fully formed around E16.5, about 3-4 days before birth (8, 11). This timing corresponds to approximately E60 in human development, with about 220 days until birth (8, 9). Genetic studies in mice, through systematic deletion of CDH-causative genes using Cre-drivers targeting critical cell lineages in diaphragm development, suggest the connective tissue lineage is a key cell population driving CDH development (8, 12–14). Complementary studies investigating the timing of gene deletion indicate that there are critical windows for when loss of CDH genes correlate with severity of herniation, with early genetic deletion leading to an absence of diaphragm muscle and late deletion leading to normal muscle development (9). The etiology and likely mechanisms underlying CDH are complex, suggesting many avenues connecting genetic aberrations to overt herniation and potential morbidity.

Similar to other structural congenital anomalies, CDH can present in many forms ranging in clinical severity. CDH can present as the sole structural defect (isolated) or can accompany defects in other organ systems in addition to lung and diaphragm (complex/syndromic). Common comorbidities include congenital heart disease (CHD) and neurodevelopmental disorders, though anomalies can involve multiple other organ systems (2). This size of the hernia is strongly associated with severity, presumably because larger diaphragm defect allows more abdominal contents to penetrate the thoracic cavity and mechanically compress the developing lungs. In addition, the location of the diaphragm defect is also important, with dorsal or posterolateral hernias associated with the more severe presentations (1, 3). Herniations located on the ventral medial edge of the diaphragm are often sub-clinical, presumably due to less impact on lung function, and can go unnoticed by clinicians or patients for many years (1, 3). The genetics underpinning both size and location of hernias remains understudied, although it has been hypothesized that somatic mosaicism may underly some forms of CDH in which a herniation occurs in one part of the diaphragm surrounded by correctly patterned muscle (6, 8). The timing of expression of genes harboring a deleterious variant may also impact the location of the defect, given what is known about diaphragm development. Lung development abnormalities independent of diaphragm defects have also been suggested to drive severity of presentation (2, 15). This has been termed the “dual-hit” hypothesis, with genetic disruption of lung development serving as the first hit and the mechanical pressure of invading abdominal contents serving as the second hit (15). This is supported by both genetic and teratogenic models of CDH (2, 15). Understanding the developmental mechanisms and genetic aberrations that lead to these different forms of CDH remains a critical goal for improving patient diagnosis and outcomes.

To extend our understanding of CDH genetics, we generated mouse models for two novel gene variants associated with CDH identified as part of the Gabriella Miller Kids First Sequencing Initiative. These variants arose in genes (*CDC42BPB* and *CNOT1)* not previously implicated in CDH development, but predicted to be causative for severe, isolated CDH. These genes were prioritized based on known gene function, expression within the developing diaphragm, and previous literature suggesting they are likely to play a critical role in diaphragm development.

*CDC42BPB* encodes the protein MRCKβ (Myotonic Dystrophy Protein Kinase-Like Beta), a serine kinase that acts downstream of the GTPase *CDC42* (16). CDC42, through MRCKβ and related MRCK family members, is an actin-myosin cytoskeletal regulator, playing a key role in cell motility across multiple cell types (16–18). CDC42BPB has been shown to directly phosphorylate key cytoskeletal components, such as myosin II regulatory light chain (MLC) (16, 19), and form complexes with other motility-related proteins at leading edges of cell migration (20). In humans, *CDC42BPB* haploinsufficiency has been associated previously with neurodevelopmental defects, specifically Chilton-Okur-Chung syndrome, an autosomal dominant condition characterized by structural brain and neurodevelopmental disorders (21).

*CNOT1* encodes a critical scaffold protein for the RNA regulation complex CCR4-NOT (Carbon Catabolite Repression 4-Negative On TATA-less). The CCR4-NOT complex, including CNOT1, is highly evolutionarily conserved (22). The protein complex has been associated with many critical functions of mRNA regulation and turnover, including transcript initiation, adenylation of mRNA, and ubiquitylation (23). Through serving as scaffolding to other enzymatic CNOT protein family members, CNOT1 has been shown to be essential for the deadenylation function of the complex and overall cell viability (24). Due to this critical role, loss of CNOT1 in the mouse is embryonic lethal (25), though tissue specific deletion of CNOT1 has been used to study its function in liver (26) and heart (25). *De novo* mutations in *CNOT1* have been associated previously with neurological developmental disorders (27–29), including holoprosencephaly, and pancreatic agenesis (30). Single nucleotide polymorphisms (SNPs) in the *CNOT1* locus have also been associated with cardiovascular disease (31, 32) and pediatric leukemia (33) through GWAS studies. These collective findings demonstrate the pleiotropic roles of both CNOT1 and CCR4-NOT in development and disease.

In this study, we demonstrate that each of these genes (*Cdc42bpb* and *Cnot1*) plays a role in diaphragm development and alterations in expression lead to different forms of CDH in mice. Both loss of *Cdc42bpb* expression and modeling of the patient missense variant leads to ventral CDH in mice through disruption in patterning of diaphragm muscle. Conversely, creating the patient variant in *Cnot1* leads to dorsal CDH with low penetrance, potentially driven by changes in transcription regulation. These findings establish that both genes are critical in diaphragm development and are causative for CDH, though the diaphragm defects differ in severity and location between mice and human.

## Results

The patient variants modeled were discovered through genome sequencing of a cohort of 827 CDH patient-parent trios (34). Both patients are male, unrelated, and presented with an isolated left, dorsal congenital diaphragmatic hernia with no history of pulmonary hypertension or pulmonary insufficiency. Both variants, c.3536C>T p.T1179M *CDC42BPB* and c.1867C>T p.R623W *CNOT1* arose *de novo* and have CADD scores of 29 and 27, respectively. The patient with the *de novo CDC42BPB* variant also showed normal neurodevelopmental outcome with a Vineland Adaptive Behavioral Scale score of 95, with assessment taking place given previous association of *CDC42BPB* damaging variants with neurodevelopmental disorders (21). The patient with the *de novo CNOT1* variant had no other reported phenotypes. Both genes have not previously been implicated in CDH or diaphragm development.

At the time of patient variant discovery, a knockout allele of *Cdc42bpb* generated by targeted deletion of the second exon (the first full coding exon) was under investigation as part of the Knockout Mouse Phenotyping Program (35) at The Jackson Laboratory, (C57BL6NJ-*Cdc42bpb^em1(IMPC)J^*/J;MGI:5812907; homozygous KO referred to as *Cdc42bpb^-/-^*; Figure 1A). Intercrosses of heterozygotes to generate homozygous cohorts revealed a high rate of pre-wean (prior to 3 weeks of life) lethality (Figure 1B), with only 2/604 weaned animals having two copies of the exon 2 deletion. We further characterized the sub-viability finding using the KOMP2 embryonic phenotyping pipeline, which includes iodine-contrast micro-computed tomography (μCT) analysis at E18.5. From initial phenotyping, *Cdc42bpb^-/-^* leads to overall smaller embryo size and pallor (Figure 1C; additional observations available on the IMPC portal: mousephenotyping.org/data/genes/MGI:2136459). Deletion of exon 2 was confirmed to lead to loss of protein expression via western blot using protein isolated from E15.5 diaphragms of wildtype and homozygous embryos (Figure 1D).

**Figure 1:**
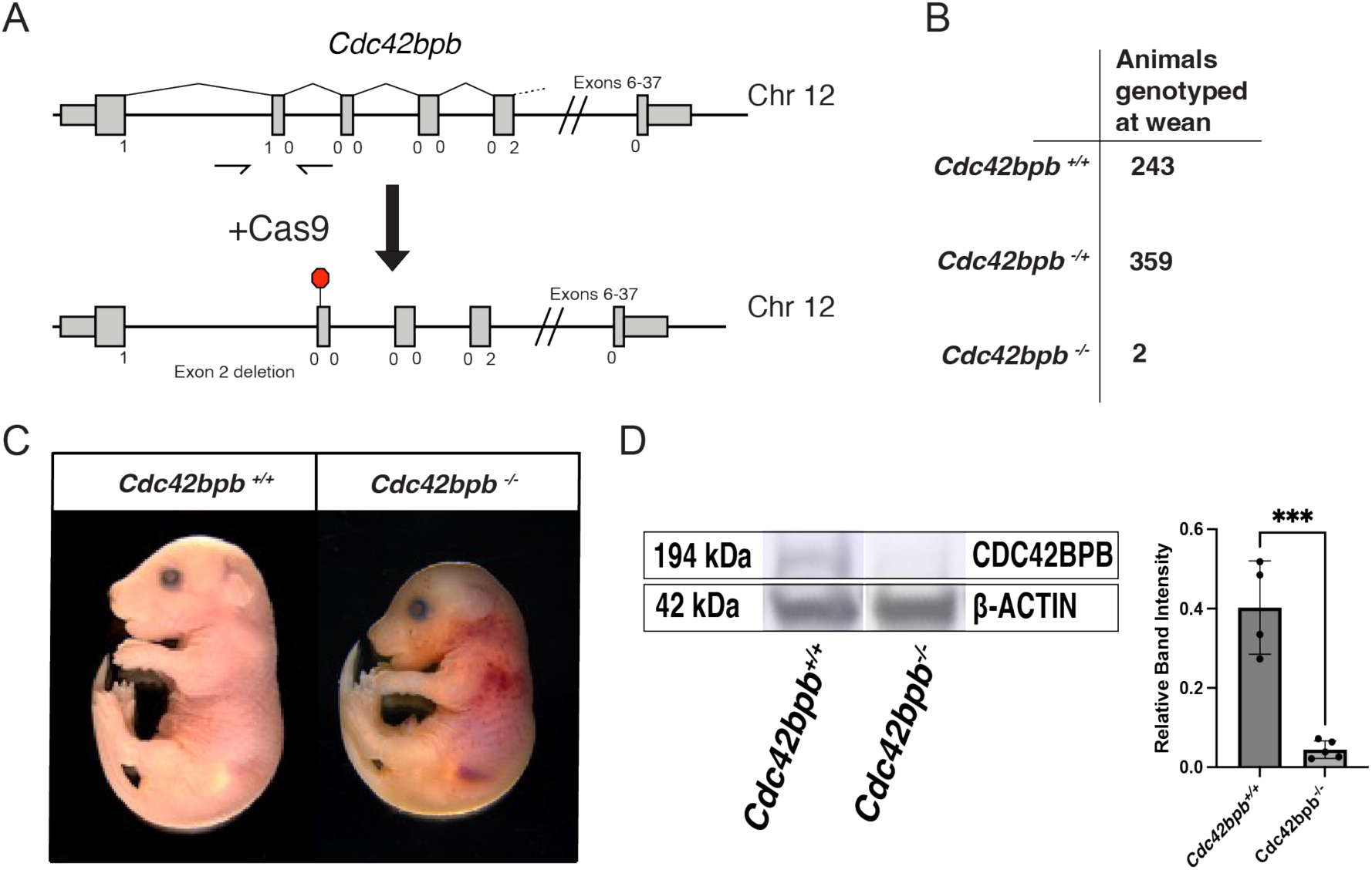
Homozygous deletion of *Cdc42bpb* exon 2 leads to pre-wean lethality. A) Targeted deletion of the second exon results in a reading frame shift and a premature stop codon at the tenth amino acid of exon 3. B) Table of genotyped mice at wean. C) Representative images of E18.5 embryos, showing smaller size and minor hemorrhage in the mutant sample. D) CDC42BPB protein expression is significantly reduced in *Cdc42bpb^-/-^* E15.5 embryonic diaphragms (N= 5) compared to wild-type controls (N=4). *** =<0.001 by Student’s t-test.

To assess for CDH, we performed a detailed assessment of internal organ structure using μCT data generated as part of the KOMP2 screen at E18.5, supplemented by scans at additional time points at E17.5 and postnatal day 1 (P1). From the cranial sections of scanned embryos, incomplete diaphragm closure and minor eventration of the liver were discovered in *Cdc42bpb^-/-^* embryos compared to *Cdc42bpb^+/+^* littermates, as highlighted by the yellow outlines in Figure 2. Three-dimensional segmentation and rendering of the developing diaphragm using ITK-SNAP (36) revealed that the herniation is consistently located ventrally, spanning from the ventral midline to the central tendon with a varying loss of muscle (Figure 2). The distance between the most ventral edges of the patterned muscle is significantly increased in *Cdc42bpb^-/-^* as compared to both *Cdc42bpb^-/+^* and *Cdc42bpb^+/+^* diaphragms (Figure 2C), demonstrating that this diaphragm phenotype is severe and highly penetrant, but one copy of *Cdc42bpb* is sufficient for normal diaphragm development. Although this does not phenocopy the human patients dorsally located diaphragmatic hernia, it does corroborate the causality of *Cdc42bpb* for CDH.

**Figure 2:**
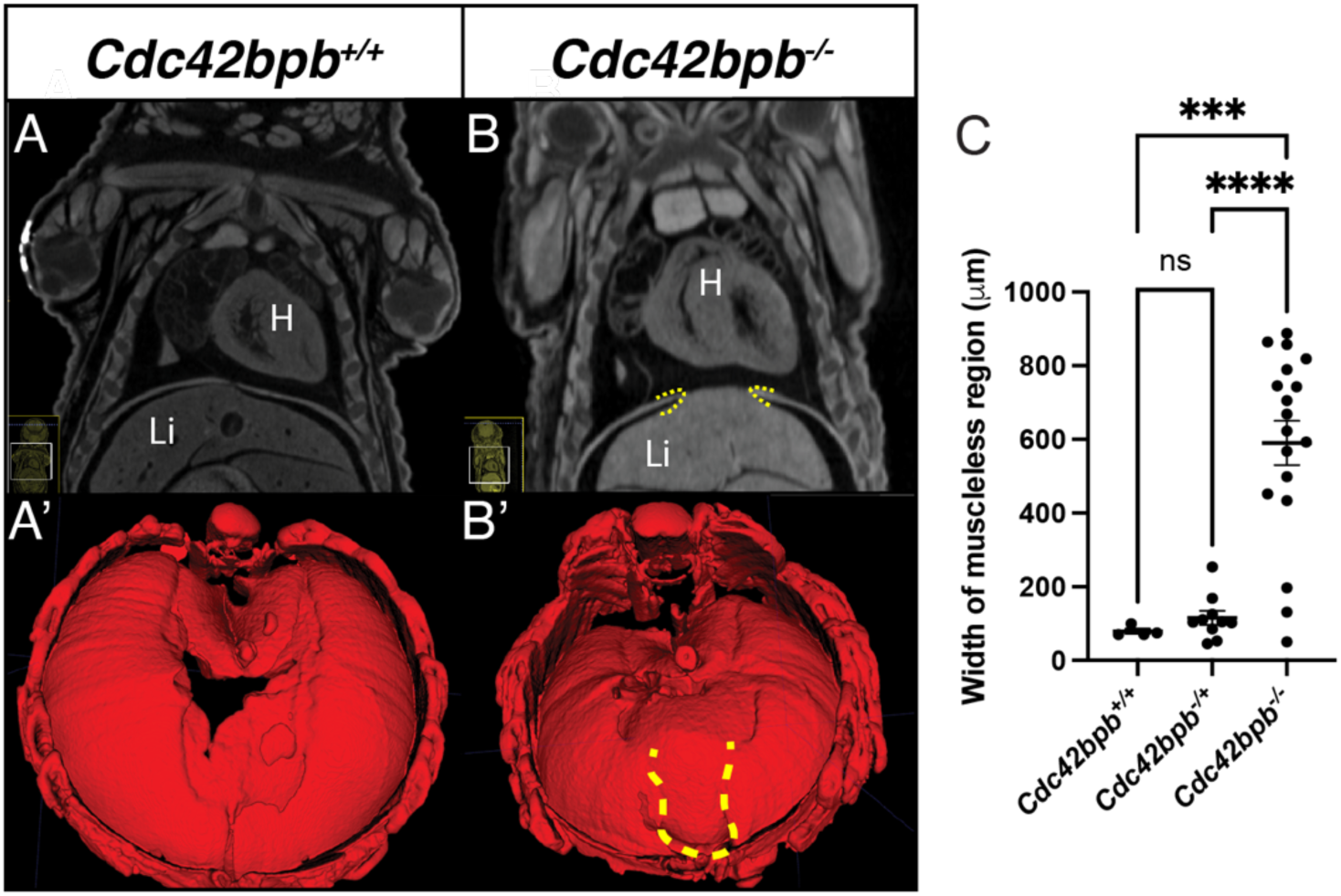
*Cdc42bpb* mutants show ventral diaphragm herniation. A) Whole body μCT scanning reveals loss of ventral diaphragm muscle (border of muscleless region highlighted with dashed-line). Under this muscleless region the liver is beginning to herniate. C) Quantitation of the distance between left and right coastal muscle sides. H-Heart, Li-Liver, ***= <0.001, ****= <0.00001 by one-way ANOVA.

The region of the diaphragm altered in *Cdc42bpb^-/-^* is the last to be patterned during development (8, 9). As the ventral midline is the last region where myofibers form (around E16.5), there is often a small “gap” at the midline between the left and right coastal muscle groups. *Cdc42bpb^-/-^* embryos consistently have a significantly greater distance between the left and right muscle groups compared with both *Cdc42bpb^-/+^* and *Cdc42bpb^+/+^* at the same embryonic stage (Figure 3). To determine whether this defect in muscle patterning is due to developmental delay, diaphragms from E14.5 to E16.5 were dissected and developing muscle fibers were visualized colorimetrically with alkaline phosphatase staining against Myosin (Skeletal, fast) protein. As early as E14.5 there is a significant distance from the ventral leading edges between the migrating developing muscle groups on the left and right side of the embryo, suggesting the muscle pattern is either delayed or altered throughout diaphragm development (Figure 3).

**Figure 3:**
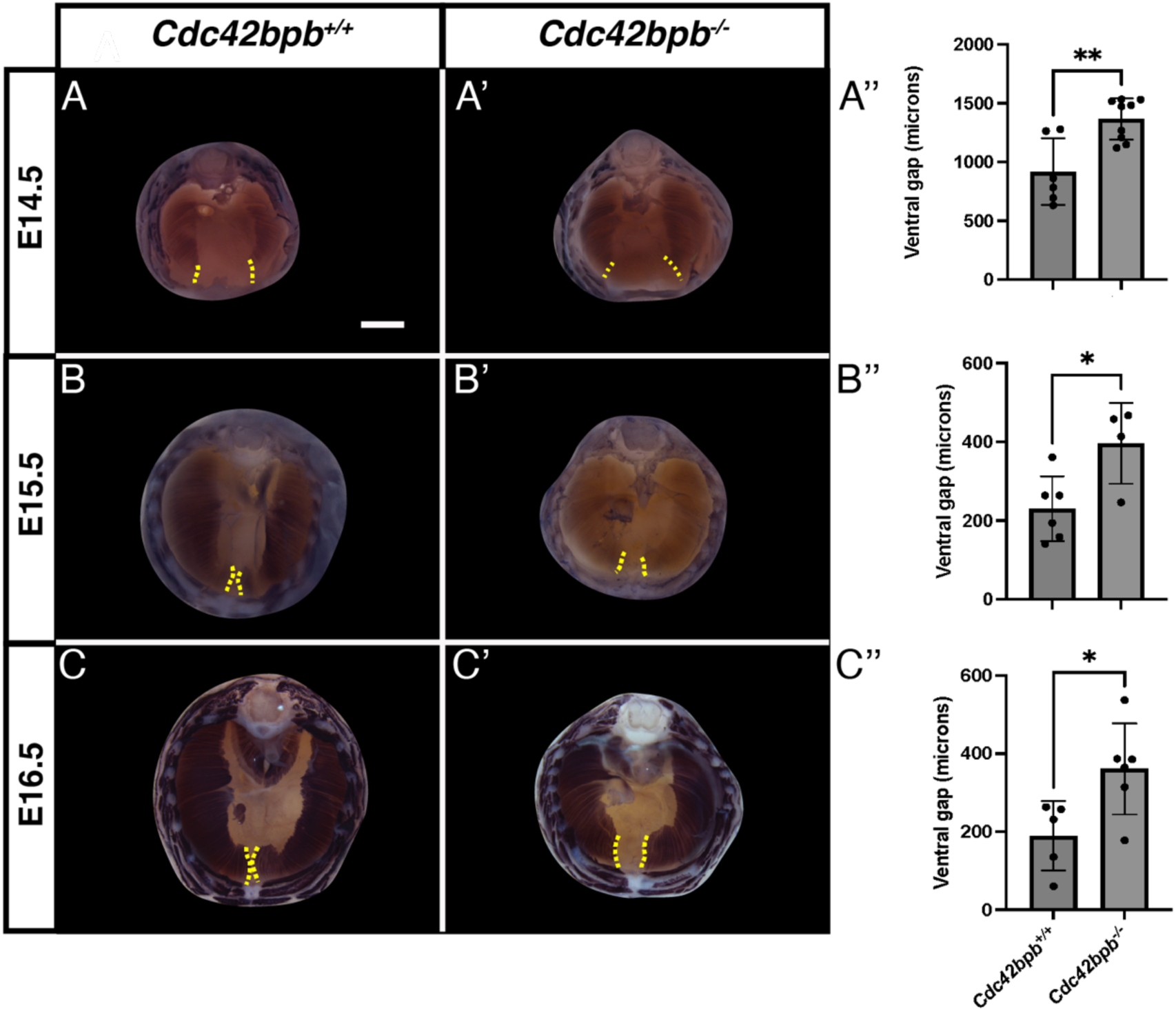
Developmental timecourse of diaphragm formation in Cdc42bpbp mutants. Dissected diaphragms stained for skeletal myosin protein (purple) at a) E14.5, b) E15.5 and c) E16.5. *Cdc42bpb^+/+^* representative images in left column, *Cdc42bpb^-/-^* representative images in right column showing differences of ventral muscle patterning and distance between left and right ventral muscle edges. Edge of patterned muscle fibers highlighted by dashed yellow line. Ventral gaps were measured across the midpoint of patterned muscle and quantified. *=<0.05, **=<0.01 by Student’s t-test. Scale bar-1000 μm.

Although highly penetrant, the *Cdc42bpb^-/-^* diaphragm ventral defect is not predicted to cause embryonic lethality, and ventral defects often present as sub-clinical in humans (3). To determine a potential driver of the neonatal lethality, a thorough manual assessment of internal structures revealed clear ventricular septation defects (VSDs) in 9/9 *Cdc42bpb^-/-^* E18.5 embryos examined (Figure 4). *Cdc42bpb^-/-^* neonates died during postnatal day 1, significantly earlier than *Cdc42bpb^-/+^* and *Cdc42bpb^+/+^* littermates. The small number of incidental deaths of *Cdc42bpb^-/+^* and *Cdc42bpb^+/+^* littermates were likely due to stress from handling (Figure 4D). Along with VSDs, lung gross morphology differences were also detected from *Cdc42bpb^-/-^* μCT scans. From 3D rendering of scanned images,16/18 lungs present with blunted distal edges and accessory lobe horn, though overall lung volume is unchanged between *Cdc42bpb^+/+^* and *Cdc42bpb^-/-^* animals (Supplemental Figure 1A-B’, yellow dashed lines). The dysmorphology of E18.5 lungs suggested minor patterning defects during mid-gestation. Bronchial development was visualized using whole mount immunostaining against cytokeratin in isolated lungs from E13.5 and E14.5. As the E18.5 phenotype appears to impact all lobes, for ease of dissection the left lobe was used for analysis at the mid-gestation timepoint (Supplemental Figure 1D and E). Counting of the distal lung buds at each stage revealed no significant differences (Supplemental Figure 1F). This suggests the apparent later stage dysmorphology may be due to a more subtle disruption of lung maturation at later time points.

**Figure 4:**
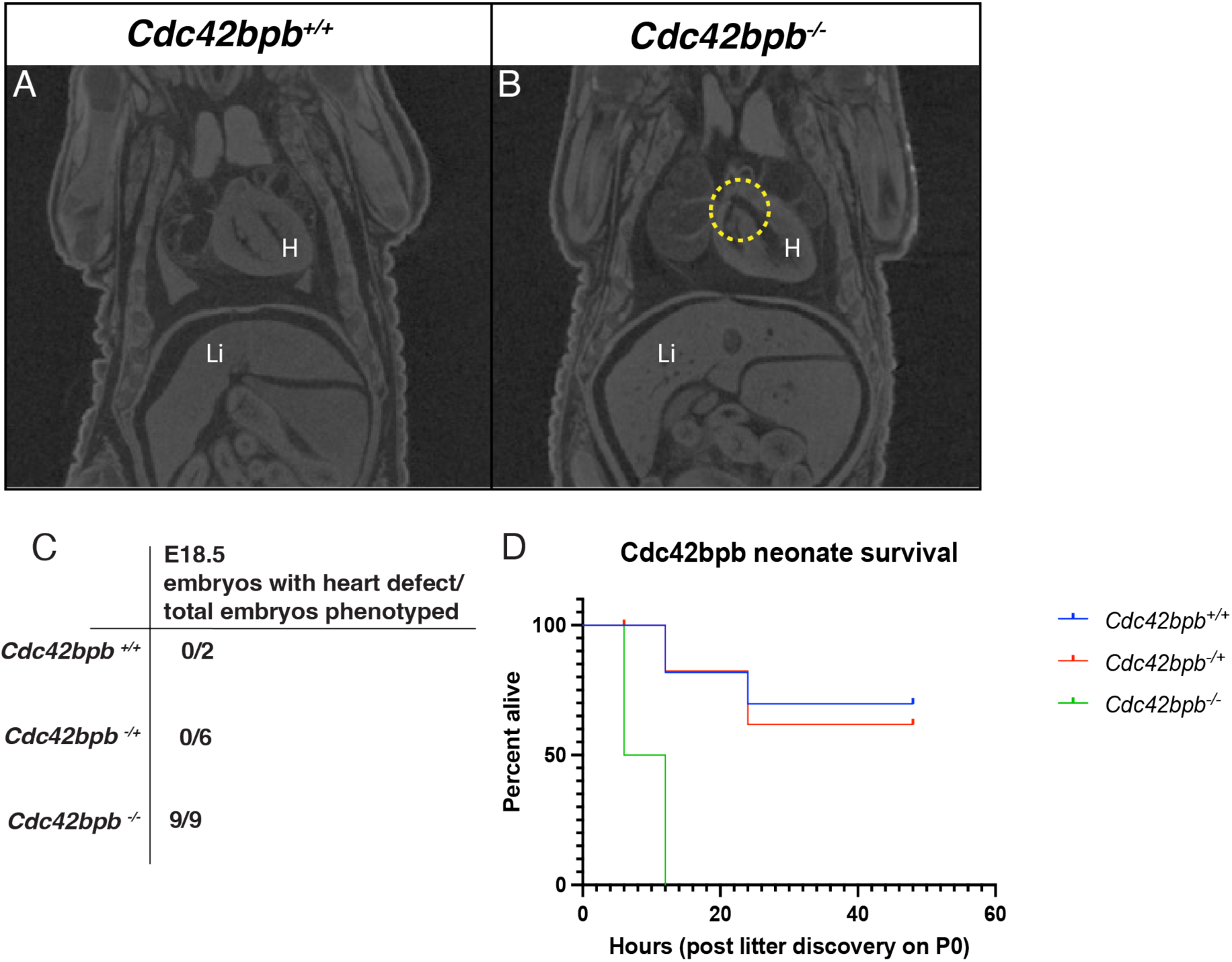
Heart defects and perinatal lethality in *Cdc42bpb^-/-^* mutants. Coronal μCT sections of A) *Cdc42bpb^+/+^* and B) *Cdc42bpb^-/-^* highlighting the septation defect (yellow dash line). C) Number of samples with observable septation defects by genotype. D) Neonate survival plot, with hour 0 set as when litter was discovered on postnatal day 0.

To determine the cell populations impacted, the lungs of E18.5 and P0 *Cdc42bpb^-/-^* were assessed for pulmonary hypoplasia phenotype which may coincide with the diaphragmatic defects. *Cdc42bpb^+/+^* and *Cdc42bpb^-/-^* mouse lungs appeared morphologically similar by hematoxylin and eosin (HCE) staining (Supplemental Figure 2A). Through staining of HOPX (Homeodomain-only Protein Homeobox), which marks the alveolar type I (ATI) cells responsible for gas exchange, E18.5 *Cdc42bpb^-/-^* mice exhibited a reduction in ATI populations (Supplemental Figure 2B-C). Despite this alveolar differentiation defect in E18.5 *Cdc42bpb^-/-^* mice, the P0 mutants did not exhibit a reduction in ATI cells and instead saw a recovery of the ATI population (Supplemental Figure 2D). Staining for SPC (Surfactant Protein-C), a marker for the alveolar type II (ATII) cells which are required for surfactant production, revealed no population differences (Supplemental Figure 2E). Transitional ATI/ATII populations were similar between control and mutant mice (Supplemental Figure 2F). Airway epithelial populations such as club cells, marked by CC10+ cells, showed no quantitative difference (Supplemental Figure 2G). The pulmonary vasculature was assessed via α-SMA and VWF staining, respectively marking the smooth muscle wall and the endothelial cells, but there was no difference in the vascular wall thickness between *Cdc42bpb^+/+^* and *Cdc42bpb^-/-^* mice (Supplemental Figure 2H). These data suggest *Cdc42bpb* plays a minor role in lung development and function, and loss of the gene does not impact respiration.

Phenotyping of the KOMP deletion mouse line demonstrates a role of *Cdc42bpb* in diaphragm development; however, the impact of the patient missense variant, C>T p.T1179M is unclear. To determine the effect of this variant, we employed CRISPR/Cas9 editing, and homology-directed repair (HDR) to create the orthologous C>T transition in mouse zygotes and directly characterized founder (F0) embryos (37, 38) (see Methods). Edited E17.5 and E18.5 embryos were harvested, genotyped, and scored for diaphragm defects using μCT whole embryo scanning (Figure 5). Genotypes were inferred from Sanger trace data using Inference of CRISPR Edits (ICE) software (ICE CRISPR Analysis. 2025. v3.0. EditCo Bio) and scored for successful installation of the target base (Knockin Score) or creation of an insertion or deletion (indel) leading to frameshift and a predicted gene knockout (Knockout Score) (See Supplemental Table 1 for ICE generated scores). Embryos with one (20-70 Knockin Score) or two (70-100 Knockin Score) copies of the patient variant presented with minimal ventral diaphragm gap distances, whereas embryos with both copies of *Cdc42bpb* predicted to be lost (70-100 Knockout Score) present with ventral gaps similar to the *Cdc42bpb^-/-^* (Figure 5). These data support the assertion that any bi-allelic loss of *Cdc42bpb* leads to a diaphragm defect but do not validate the patient variant as directly causative for CDH.

**Figure 5:**
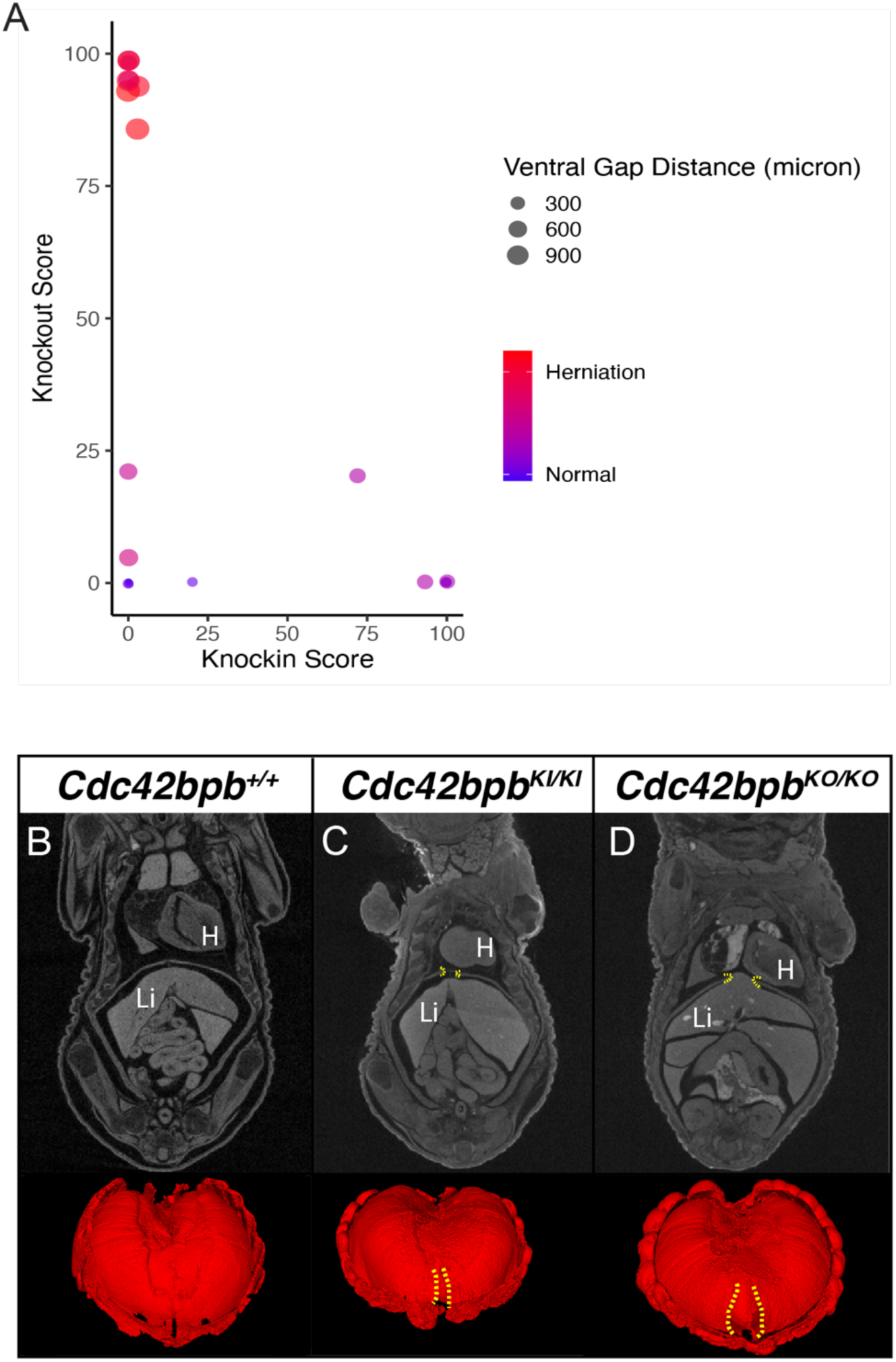
Functional validation of the *CDC42BPB*-T1179M variant for CDH in F0 mouse embryos. A) Scatterplot comparing ICE (see methods) knockin (specific variant; x-axis) and knockout (frame-shifting indel; y-axis) scores for each scanned embryo (datapoints). Herniation status of E18.5 embryo denoted by datapoint color, and ventral gap distance denoted by size of datapoint. B), C), and D) representative coronal sections and diaphragm renders for unedited, 100% knockin and 100% knockout embryos, respectively. Ventral gaps in both section and render highlighted by yellow dash lines. H-Heart, Li-Liver.

The second candidate variant in this study was a predicted damaging missense variant in *CNOT1*, a critical gene in the CCR4-NOT complex not previously associated with CDH*. Cnot1* is a cell essential gene, with complete loss leading to embryonic lethality in mice (25). To determine whether the R623W mutation causes CDH, we engineered the orthologous C>T variant using CRISPR/Cas9 mediated editing and established a germline model, C57BL/6NJ-*Cnot1^em1Murr^*/Murr (referred to as *Cnot1^R623W^*; see Figure 6A and Methods). Both heterozygous and homozygous animals are viable, are fertile and show no significant change in expected Mendelian ratios of genotypes (*Cnot1^+/+^* 77/351, *Cnot1^-/+^* 189/351 and *Cnot1^-/-^* 85/351). Late-stage embryos and early-life neonates (E18.5-P2) were collected and either μCT imaged or dissected to score diaphragm defects (Figure 6B-E).

**Figure 6:**
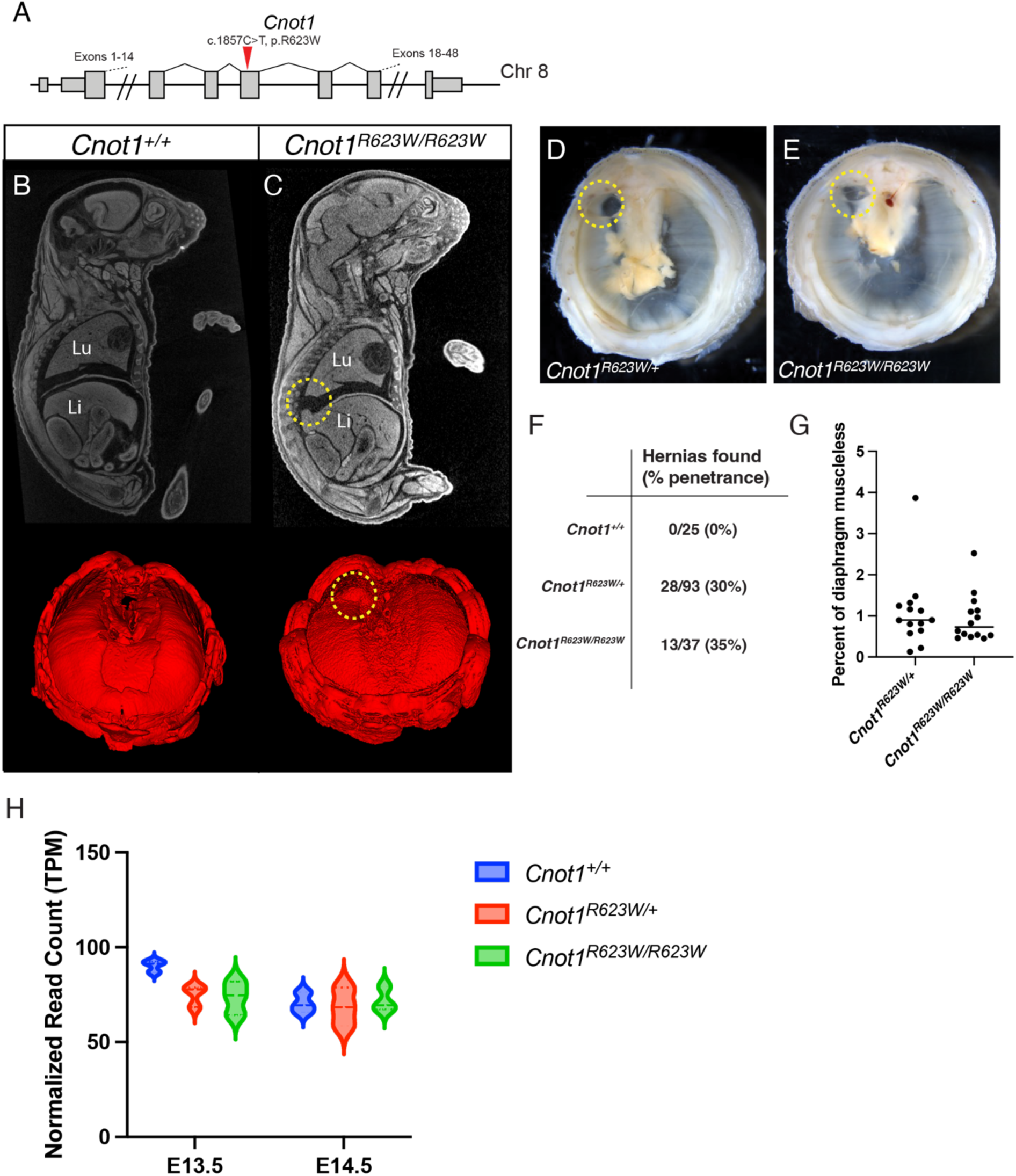
The *CNOT1*-R623W patient-specific variant leads to dorsal diaphragmatic hernia with incomplete penetrance in both heterozygous and homozygous mouse embryos. A) Location of the patient-specific missense variant inserted into exon 17 of mouse *Cnot1*. Representative images of B) *Cnot1^+/+^* and C) *Cnot1^RC23W/RC23W^* of μCT scanned E18.5 embryos and rendered diaphragms from scans. Dorsal diaphragmatic hernia in *Cnot1^RC23W/RC23W^* highlighted by yellow dash lines. Dissected P1 diaphragms, with livers removed, with yellow dashed lines highlighting dorsal herniations present in D) *Cnot1^+/R623W^* and E) *Cnot1^RC23W/RC23W^*. Total number of diaphragmatic hernias discovered in E17.5-P1 animals across all genotypes, with percent penetrance noted. G) Quantitation of muscleless regions (area of herniated diaphragm/area of total diaphragm muscle) between *Cnot1^+/R623W^* and *Cnot1^RC23W/RC23W^*. Not significant by Student’s t-test. H) RNA expression of *Cnot1* from bulk RNA sequencing from E13.5 and E14.5 diaphragms (N=3 per genotype). No significant difference was found in normalized read count (Transcript per Million or TPM) by one-way ANOVA.

Dorsally located amuscular regions were discovered in animals with both one or two copies of the R623W allele, at similar low penetrance (Figure 6F). The relative size of the defect is also similar between heterozygous and homozygous animals that present with a diaphragm defect (Figure 6G). From bulk RNA-sequencing of E13.5 and E14.5 diaphragms no significant change in expression between genotypes were found (Figure 6H, N=3 diaphragms per genotype per stage). Compared to the ventrally located defect found in the *Cdc42bpb^-/-^*, the *Cnot1^R623W^* phenocopies the dorsally located diaphragm defect reported in the human patient. As found in human patients (1, 3), these dorsally located hernias are more likely to lead to a compression of the developing lungs, as seen in the highlighted gene of the μCT sagittal section in Figure 6C. While phenocopying the patient form of CDH, the diaphragm defect appears sublethal due to low penetrance, given the minimal impact on lethality at wean.

*Cnot1* encodes a critical component of the CCR4-NOT protein complex, which regulates mRNA processing by deadenylation of the poly(A) tail (22, 24), and thus we hypothesized that the patient’s variant impacts complex function and global gene expression. To test this, developing diaphragms were collected at E13.5 and E14.5 from *Cnot1^+/R623W^*, *Cnot1^RC23W/RC23W^* and wildtype littermates, total RNA was isolated and RNA sequenced (see Methods). At E13.5 gene expression was not significantly changed when comparing wildtype to *Cnot1^+/R623W^* or *Cnot1^RC23W/RC23W^* samples (Supplemental Figure 2, Supplemental Table 2). At E14.5 a minimal number of genes are differential expressed comparing either *Cnot1^+/R623W^* or *Cnot1^RC23W/RC23W^* to wildtype (Figure 7A). When analyzed with STRING (39) an interaction network forms between genes differentially express in *Cnot1^+/R623W^* compared to *Cnot1^+/+^* (Figure 7B and C), and the differentially expressed genes are enriched for “abnormal morphology” and “lethality” Gene Ontology (GO) Mammalian Phenotype Ontology (MPO) (40) terms (Supplemental Table 3) from the Mouse Genome Database (MGD) (41). When comparing *Cnot1^RC23W/RC23W^* and wildtype, only 7 genes were found to be differentially expressed, and no enrichment terms were found when analyzed with STRING.

**Figure 7:**
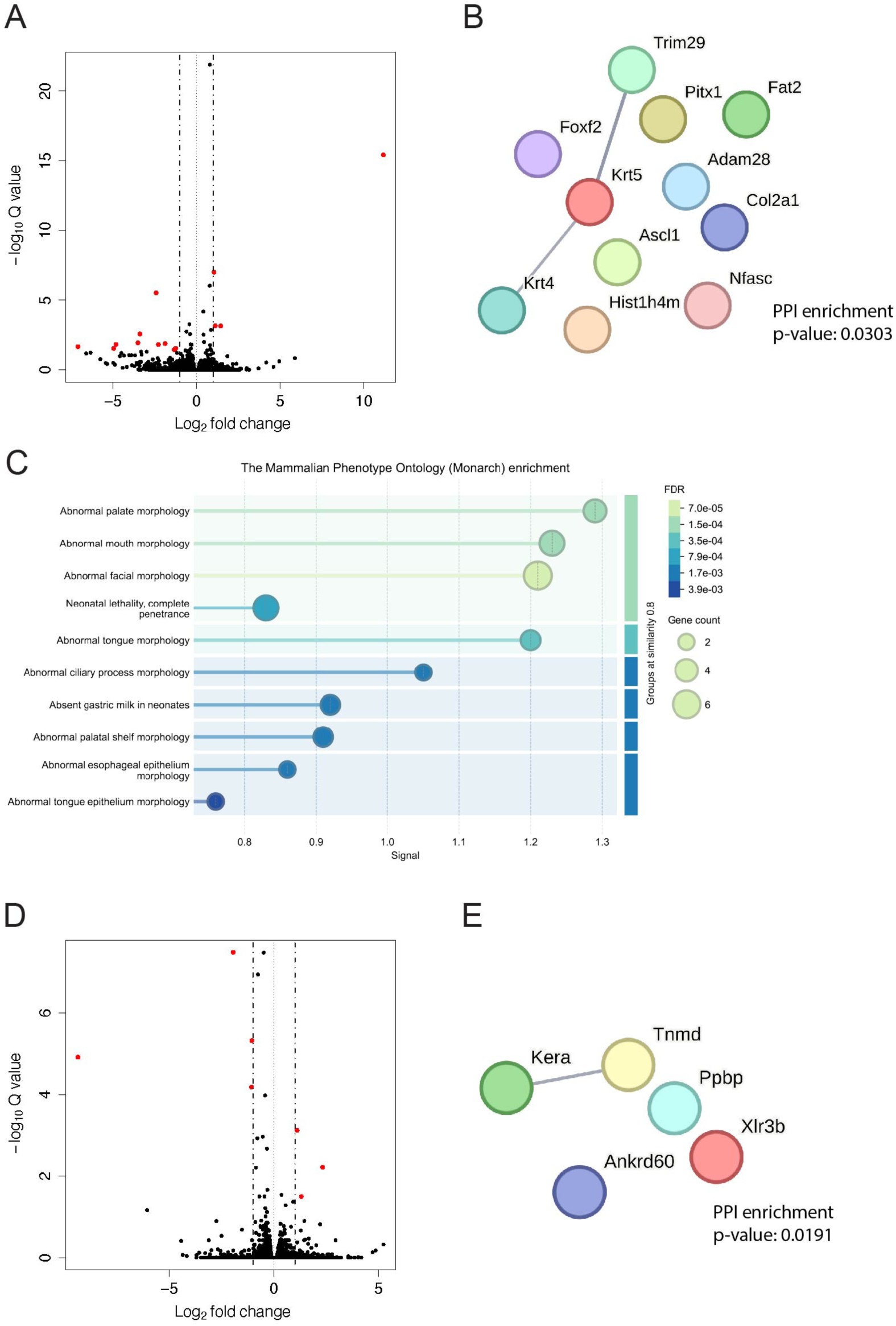
Transcriptional changes in Cnot1 mutant diaphragms at E14.5. Volcano plots highlighting differential expressed genes in comparisons of: A*) Cnot1^+/+^* vs. *Cnot1^+/R623W^* and B) *Cnot1^+/+^* vs *Cnot1^RC23W/RC23W^*. Red datapoints represent genes with absolute log_2_ fold change greater than 1 and p-adjusted (Q) value less than or equal to 0.05. STRING network analysis of significant differentially expressed genes for C) *Cnot1^+/+^* vs. *Cnot1^+/R623W^* and D) *Cnot1^+/+^* vs *Cnot1^RC23W/RC23W^*. Lines represent genes with known interaction. E) STRING ontology enrichment of *Cnot1^+/+^* vs. *Cnot1^+/RC23W^* differentially expressed genes, highlighting Monarch terms significantly enriched. No enrichment terms were found for *Cnot1^+/+^* vs *Cnot1^RC23W/RC23W^*.

Given that *Cnot1* is critical in post-transcriptional mRNA processing, we also investigated changes in transcript isoforms in samples with the *Cnot1^R623W^*. Comparison of either *Cnot1^+/R623W^* or *Cnot1^RC23W/RC23W^* transcript isoform expression levels to *Cnot1^+/+^* revealed many differentially expressed transcripts (DETs) at both E13.5 (Supplemental Figure 4) and E14.5 (Supplemental Figure 5, Supplemental Table 4). STRING network analysis revealed an enrichment of related DETs only at E14.5 comparisons, with protein-protein interaction (PPI) enrichment p-values of 0.00241 for *Cnot1^+/+^* x *Cnot1^+/R623W^* and 0.0242 for *Cnot1^+/+^* x *Cnot1^RC23W/RC23W^* compared to p-values of 0.496 and 0.107 for equivalent comparisons at E13.5. From functional enrichment term analysis of DETs, many terms across STRING databases were found (Supplemental Table 5), varying across each comparison. Across all comparisons only the UniProt annotated keywords “Alternative splicing” and “Phosphoprotein” were consistently significantly enriched (Supplemental Figures 4C-F and 5C-F). “Alternative splicing” is likely in part due to the comparison of distinct gene isoforms (created by mRNA splicing) being compared. An enrichment of phosphorylated proteins may be more biologically relevant, though this keyword encompasses many proteins in the mouse genome (7592 proteins within the STRING Uniprot database). Analyzing significant DETs shared across the two comparisons at each stage (17 DETs at E13.5 and 22 at E14.5, Figures 8 and 9) did reveal the “Phosphoprotein” keyword remains enriched at E13.5 (with 14/17 proteins tagged, Figure 8D red highlighted proteins), though only “Bromo domain” (3/22 proteins) remains at E14.5 (Figure 9D, red highlighted proteins).

**Figure 8:**
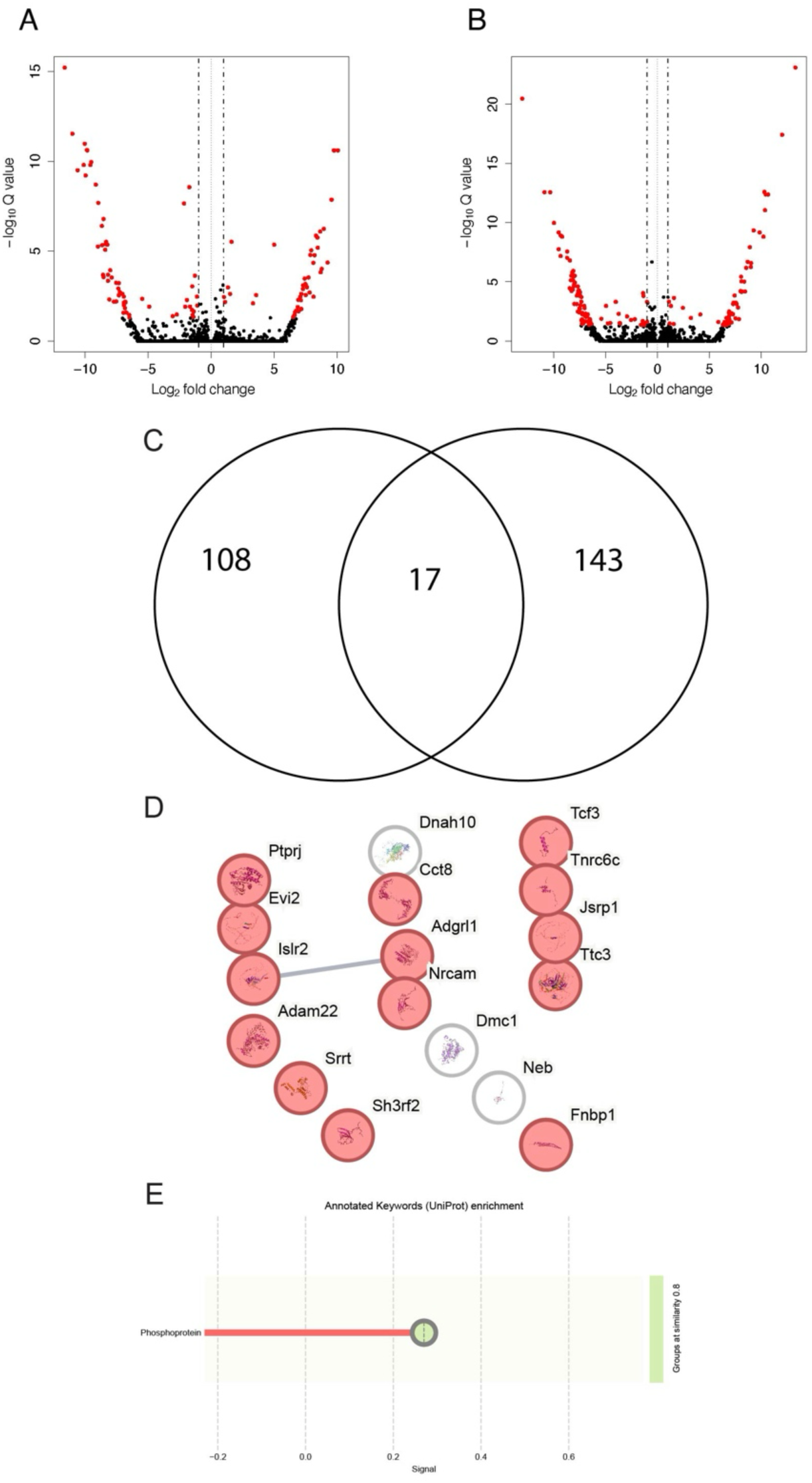
Differentially expressed transcription isoforms in *Cnot1* mutants at E13.5. Volcano plots of transcript differential analysis between A) *Cnot1^+/+^* vs. *Cnot1^+/R623W^* and B) *Cnot1^+/+^* vs. *Cnot1^RC23W/RC23W^*. Transcripts (datapoints) with absolute log_2_ fold change greater than 1 and p-adjusted (Q) values less than 0.05 highlighted in red. C) Number of significantly expressed transcripts unique and shared between *Cnot1^+/+^* vs. *Cnot1^+/R623W^* and *Cnot1^+/+^* vs. *Cnot1^RC23W/RC23W^* comparisons. D) STRING analysis on transcripts shared between differential comparisons. E) STRING ontology analysis enriched term within shared significant transcripts. Genes within the enriched term highlighted in red in D).

**Figure 9:**
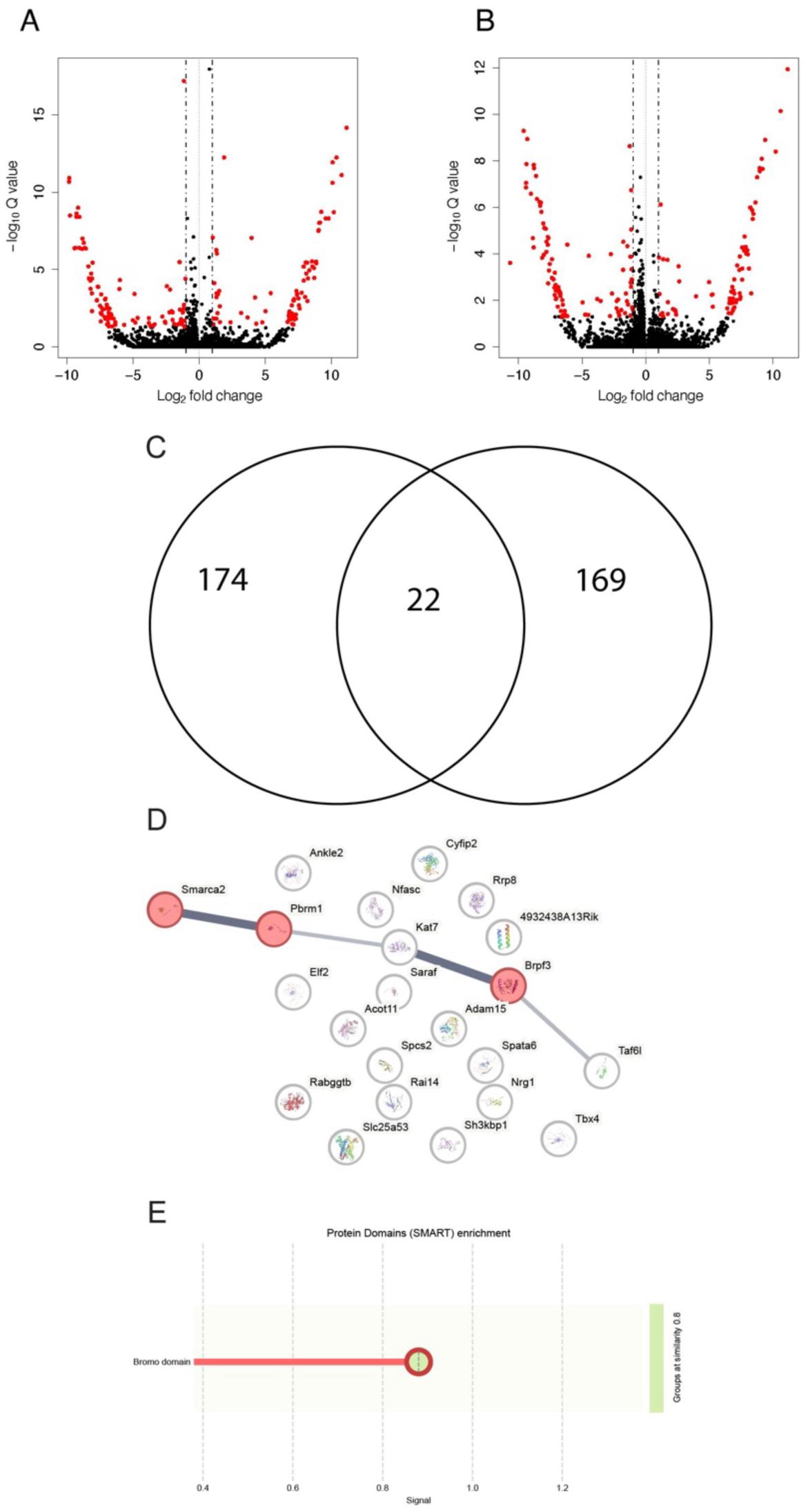
Differentially expressed transcription isoforms shared in in *Cnot1* mutants at E14.5. Volcano plots of transcript differential analysis between A) *Cnot1^+/+^* vs. *Cnot1^+/R623W^* and B) *Cnot1^+/+^* vs. *Cnot1^RC23W/RC23W^*. Transcripts (datapoints) with absolute log_2_ fold change greater than 1 and p-adjusted values less than 0.05 highlighted in red. C) Number of significantly expressed transcripts unique and shared between *Cnot1^+/+^* vs. *Cnot1^+/R623W^* and *Cnot1^+/+^* vs. *Cnot1^RC23W/RC23W^* comparisons. D) STRING analysis on transcripts shared between differential comparisons. E) STRING ontology analysis enriched term within shared significant transcripts. Genes within the enriched term highlighted in red in D).

Conditional loss of *Cnot1* has been shown to lead to global changes in polyadenylation, and the patient predicted damaging variant is located near the region where CNOT7 and CNOT8, key proteins in the adenylation regulation, bind (22, 26). To determine if a change in polyadenylation was captured in the total RNA sequenced diaphragm samples, the tool APAlyzer (42) was used. APAlyzer uses reference polyadenylation sites (PAS) databases to identify all polyA sites across the region, then compares read density across mutant and control samples at those sites for relative expression scores, allowing for differential polyadenylation analysis at the gene level (42). Though APAlyzer found genes with signatures of polyadenylation change when wildtype samples were compared to *Cnot1^+/R623W^* and *Cnot1^RC23W/RC23W^* samples at both developmental stages (Supplemental Figure 6A-D), no gene reached significance after multiple testing correction (Supplemental Figure 6A’-D’, Supplemental Table 6). Visual inspection of polyadenylation sites of the top ten genes identified to have altered polyadenylation were found to have minimal to no change in read density across samples in the Integrative Genome Viewer (IGV) (43). Overall, there appears to be little to no detectable change in polyadenylation, although due to the low penetrance of the CDH phenotypes and potential hypomorphic nature of the *Cnot1^R623W^* allele, any change in polyadenylation or overall gene expression may be difficult to detect.

## Discussion

As clinical genomic sequencing efforts and technologies expand, the number of novel variants and genes predicted to underly disease including congenital anomalies will continue to increase. In parallel, methods to validate a gene or variant’s role in disease pathogenesis are vital. Here we present *in vivo* characterization validating the role of two novel CDH variants predicted to drive severe forms of CDH in human patients. Our models were useful for interrogating the roles of both the gene and the patient-specific variant identified. For *Cdc42bpb,* we leveraged an existing knockout resource to establish that the gene is necessary for diaphragm development. In contrast, through F0 modeling via direct phenotyping of Cas9-edited embryos we were not able to validate the causality of the patient-specific missense variant, but did further corroborate that loss-of-function mutations could result in CDH. The consistent ventral hernia phenotype seen in mice does not phenocopy the dorsal hernia seen in patients but does support a role for the gene in normal diaphragm development. Further work is needed to understand this discrepancy, which has been found in other mouse CDH models (44), and how much this phenomenon reflects species-specific differences at the genetic or developmental level. For *Cnot1,* our patient-specific missense allele resulted in dorsal, isolated CDH in both heterozygous and homozygous samples, phenocopying the patient at both zygosities, consistent with the *de novo* mutation effect. With these two mouse lines we can model two ends of the spectrum of CDH severity impacted by diaphragm defect size.

The *Cnot1^R623W^* allele leads to dorsal diaphragm hernias and has a minor impact on RNA transcription. Though phenocopying the Bochdalek (dorsal) hernia found in the human patient carrying this allele, the missense mutation did not lead to detectable lethality in either heterozygous or homozygous mice. The incomplete penetrance may lead to rare instances of lethality which were not detected in our analysis, or alternatively the herniation is not severe enough to impact animal fitness. To date no adult mouse with either one or two copies of the *Cnot1^R623W^* have been found to have a diaphragmatic hernia similar to those found in embryos or neonates. However, our findings support a role for *Cnot1* in diaphragm development and corroborate the patient missense mutation as likely contributing to CDH. Bulk RNA sequencing analysis showed two recurrent GO enrichment terms: “Phosphoprotein” and “Alternative Splicing”, the latter consistent with a splicing role for the complex. Although significant across all four comparisons, both terms have a lower enrichment effect, possibly due to the large number of genes in the “Phosphoprotein” and “Alternative Splicing” networks in the Uniprot database (45). Previously it has been shown that depletion of CNOT1 *in vitro* correlates with altered phosphorylation of critical cell cycle components leading to senescence (46). Recently, it has also been shown that phosphorylation changes of CCR4-NOT binding partners alter the amount of deadenylation of target mRNAs (47). It is likely the *Cnot1^R623W^* allele is hypomorphic, leading to more subtle changes compared to full depletion of the *Cnot1*. Although we hypothesized the deadenylation function may be altered enough in the developing diaphragm to lead to herniation, as this is a well-documented and critical function of the CCR4-NOT complex (23, 24), we did not detect a significant impact on mRNA 3’ adenylation in diaphragms with one or two copies of *Cnot1^R623W^*. This leaves at least three possibilities to be tested: 1) we are underpowered to detect a low level of APA differences in discrete transcripts in the diaphragm, consistent with variable severity and incomplete penetrance in *Cnot1^R623W^* mutants; 2) the threshold of altered adenylation in the diaphragm is so sensitive the stochastic differences across animals is enough for some to develop hernias, for which again we are underpowered to detect; or 3) the herniation associated with *Cnot1^R623W^* is not driven by APA changes in CCR4-NOT complex function. Further investigation to distinguish these mechanisms is necessary. Given that the germline knockout of *Cnot1* is embryonic lethal, conditional analysis in diaphragm-specific cell types will be critical for future mechanistic studies into the role of *Cnot1* in diaphragm development.

Homozygous loss of *Cdc42bpb* leads to large ventral gaps in the diaphragm with high penetrance, minor changes in lung morphology, and lethal heart septation defects. Similar ventral defects have been reported in other mouse models of cytoskeletal genes including a *Pls3* missense mutation and a *Myh10* knockout (44, 48). However, in both *Pls3* and *Myh10* models, ventral body wall defects were reported, which to date have not been found in *Cdc42bpb^-/-^* mice. Ventral diaphragm muscle mis-patterning was also seen when *HGF*/*MET* signaling is disrupted, both genetically and pharmacologically, in the mouse embryonic diaphragm (14). The *HGF/MET* signaling pathway is critical for muscle recruitment from the somites to the PPFs, and the authors demonstrated a loss of myogenic cells at the leading edge of muscle patterning at E15.5. This could suggest a similar loss of myogenic cells in *Cdc42bpb^-/-^* diaphragms, leading to a similar ventral gap. To date no *Cdc42bpb^-/-^* neonate has any noted respiration phenotypes, though the observation window is small, thus it is likely these noted phenotypes at E18.5 are not impacting overall lung function. However, given the difference in diaphragm phenotype between the human patient and *Cdc42bpb^-/-^* embryos gene disrupting variants in human *CDC42BPB* may lead to more severe lung defects. *Cdc42bpb^-/-^* neonate lethality is likely driven by the structural heart defects found through μCT imaging (Figure 4) and not by defects in diaphragm or lung. This co-morbidity also reflects the common presentation of CHD with CDH in human patients (1), and cardiovascular defects in patients with *CDC42BPB* variants have been reported (21). This pleiotropy across heart, lung and diaphragm is found across many CDH-related genes, both in human patients presenting with CDH, CHD or both and from *in vivo* animal modeling (1, 15). Other pleiotropic CDH genes such as *Gata4* and *Zfpm2* lead to early embryonic lethality when deleted in the mouse germline (49–51), before diaphragm development begins, unlike *Cdc42bpb* deletion leading to a cardiac septation defect impacting neonate fitness, but not early embryonic development. Further investigation into the role of *Cdc42bpb* in development across heart, lung, and diaphragm could improve our understanding of key gene pathways, cell-cell communication, and protein dynamics that lead to the co-morbidities found across all three tissues.

Presented are two novel CDH mouse models selected to determine whether the two genes likely contributed to severe, isolated, dorsal CDH in two unrelated patients. Both mouse models present with CDH, though as shown, differ in both location and phenotypic severity. This key difference appears to be due to differences in underlying cellular mechanisms, as loss of *Cdc42bpb* appears to lead to a delay and failure of the diaphragm muscle to reach the ventral edge of the diaphragm, whereas the *Cnot1^R623W^* allele leads to a subtle change in transcription. Though demonstration of the orthologous patient variant leading to CDH is crucial for validation of human genetic studies, further mechanistic investigation into the role of *Cnot1* in the diaphragm will be difficult. Outstanding questions remain for both genes, including which cell populations are affected, whether gene expression is required throughout diaphragm development, and if the gene plays similar roles in other organ systems. Similar questions remain for many other CDH candidate genes (1, 5, 6). Groups have begun to explore diaphragm gene regulation at the single cell level early in development (52) and after diaphragm development is complete (53). Continued exploration of cell population dynamics across diaphragm development is crucial in understanding the gene presented here and future validated CDH gene roles in CDH. Combining insights gained from initial validating screens, such as the F0 platform used for the *Cdc42bpb* patient mutation, ‘omics approaches to understand how the gene functions during diaphragm development, and targeted genetic manipulation of the gene. All remain critical in understanding both CDH and congenital anomalies pathogenesis, and translating these findings into improved clinical diagnostics.

## Materials and Methods

### Mice

All animal studies were performed in accordance with the Guide to the Care and Use of Laboratory Animals and were reviewed and approved by the Institutional Animal Care and Use Committee of The Jackson Laboratory.

### Model engineering

The *Cdc42bpb^em1(IMPC)J^* allele was created at The Jackson Laboratory as part of the Knockout Mouse Phenotyping Program (KOMP2) (35). Guide RNA sequences and allele design details can be found at https://www.mousephenotype.org/data/alleles/MGI:2136459/em1(IMPC)J?alleleSymbol=em1(IMPC)J#mice. Mice were maintained on a C57BL/6NJ (The Jackson Laboratory, strain #005304) background.

The *Cnot1^em1Murr^* allele was generated using CRISPR/Cas9 endonuclease-mediated genome editing. Guide RNA (CAAGTCCACCCAAAATAGAA) was selected to cleave DNA within exon 16 of the Cnot1 gene and a single stranded oligo donor with a missense R623W (AGG>TGG) and silent C624C (TGC>TGT) variant were introduced to facilitate HDR-mediated genome editing. The guide and single stranded donor were introduced to single cell C57BL/6NJ zygotes and transferred to pseudo pregnant females. Progeny were screened by DNA sequencing to identify correctly targeted pups, which were then bred to C57BL/6NJ (Stock No. 05304) mice for germline transmission. The C57BL/6NJ-*^Cnot1em1Murr^*/Murr colony was backcrossed to C57BL/6NJ for at least two generations. See Supplemental Table 7 for oligonucleotide sequences used.

### Embryo and neonate collection

Mouse embryos and neonate pups were collected from heterozygous-by-heterozygous timed matings for both *Cdc42bpb* and *Cnot1* lines. DNA was extracted from either yolk sacs (embryos) or tail tips (neonate pups) for genotyping by melt curve analysis for *Cnot1^R623W^* or standard PCR for *Cdc42bpb^-^* (see primers in supplemental table 7). Embryos were harvested through microdissection, imaged for gross morphological phenotypes on a M125 Leica dissecting microscope, crown-rump length (CRL) measurements for stage confirmation were taken, notable external phenotypes were noted and embryos/neonates were either: exsanguinated in 1x PBS with 0.2 mg/mL heparin (for E16.5-18.5) then fixed overnight in 4% paraformaldehyde (PFA) at 4°C; directly fixed overnight if younger; or, neonates were euthanized by decapitation, then fixed in 4% PFA. To score herniations diaphragms were micro-dissected after fixation as previously described (8), for *Cnot1* diaphragms liver was also removed to more easily visualize diaphragm abnormalities. To quantify *Cnot1* herniation size the total area of the herniation was divided by the total area of the diaphragm muscle from M1654 Leica dissecting microscope images. Herniation percentages between *Cnot1^+/RC23W^* and *Cnot1^RC23W/RC23W^* were statistically compared using Student’s T test with GraphPad Prism version 10.0.0 for Windows, GraphPad Software, Boston, Massachusetts USA, www.graphpad.com.

### Cdc42bpb^-/-^ neonate survival screen

Expected birth date was determined based on seminal plug in *Cdc42bpb^-/+^* timed matings, and boxes were checked every two hours throughout the day. Once pups were born and discovered CRL and body weight were measured and toe clips were taken to both identify each pup and genotype each pup using PCR amplification. Littters were checked every two hours after initial discovery and measurements were repeated every four hours.

### μCT embryo preparation, scanning and analysis

After 4% PFA overnight fixation samples were washed in PBS and placed in STABILITY buffer (35) for 5-7 days. at 4°C. Embryos were then incubated at 37°C for 4 hours to polymerize the STABILITY buffer, which was then cleaned off the sample. Embryos were then placed in 50 mL iodine-based Lugol’s Solution and incubated with agitation at room temperature for 5-7 days. Counterstained samples were then mounted in 1% agarose in 5 mL flat bottom conical tubes, stacking two samples vertical. Mounted, counterstained samples were imaged on either a Bruker Skyscan 1172 or 1272 μCT (Bruker Biospin), with 0.5 millimeter filter, 2048×2048 resolution, and pixel size of 11 μm (parameters adapted from (35)). Scan image sets were then reconstructed using NRecon software (Bruker), exporting data as transaxial bmp image stacks, which were then converted into. nrrd three-dimensional image files using Fiji (ImageJ). ITK-SNAP (36) software was used to open .nrrd files, score internal structural tissue defects and render diaphragm and lungs for further phenotyping. Lung relative volumes were collected by dividing rendered lung voxel volumes from total rendered embryo/neonate voxel volumes in ITK-SNAP. For *Cdc42bpb* rendered diaphragms, ventral gap distance measurements were taken by exporting diaphragm 3D renders to 3DSlicer and using the measure tool across the two edges of muscle at the midpoint between the ventral body wall and the central tendon. Heart defects were scored by comparing coronal and transverse sections of *Cdc42bpb^-/+^* and *Cdc42bpb^-/-^* μCT scanned hearts to *Cdc42bpb^+/+^* scanned litter mates. Kaplan-Meier survival plot and analysis was performed using GraphPad Prism version 10.0.0 for Windows, GraphPad Software, Boston, Massachusetts USA, www.graphpad.com.

### Patient variant modeling using F0 embryo CRSIPR/CasS editing

CRISPR gRNA targeting the mouse orthologous *Cdc42bpb* CDH patient variant location and single strand DNA carrying both patient missense variant and gRNA-disrupting base pair change were designed using design tools built-in to the Benchling platform (benchling.com, (54, 55)). CRISPR/Cas9 ribonucleotide complexing, E0.5 electroporation of editing components and implantation were carried out using previously described methods (56). CRISPR/Cas9 components were ordered from Integrated DNA Technologies (IDT) from the Alt-R Cas9 platform. E18.5 embryos were harvested from pseudopregnant females implanted with edited fertilized zygotes, yolk sacs were collected for DNA, and embryos were fixed and processed for μCT scanning using the same methods as embryos collected from timed matings. Embryos were scanned, scored for structural defects and diaphragms were rendered with ITK-SNAP (36). Diaphragm ventral gaps were measured using the same methods in 3DSlicer as timed matings. PCR amplified DNA was Sanger sequenced and trace files were deconvoluted using ICE (ICE CRISPR Analysis. 2025. v3.0. EditCo Bio) and percentages of variant knock-in and knockout were determined.

### Western blotting

E15.5 diaphragms were microdissected from harvested *Cdc42bpb^-/+^* x *Cdc42bpb^-/+^* as previously described (57), snap frozen in liquid nitrogen and stored at -80°C. Once embryos were genotyped through standard PCR, like genotypes were pooled to three diaphragms per tube (e.g. per sample [N] in final quantification), then lysed using RIPA buffer (Thermo Fisher Scientific) with cOmplete^TM^ Protease Inhibitor (Sigma-Aldrich) and mechanically disrupted with a disposable plastic pestle (Thermo Fisher Scientific). Protein supernatant was separated by centrifugation at 14,000rpm for 20 minutes at 4°C. Protein concentration was determined with Pierce^TM^ BCA protein assay kit (Thermo Fisher Scientific). Protein was diluted to a concentration of 1 μg/μL in RIPA and 4x Laemmli Sample Buffer (Bio-Rad) and 30 μL per sample was loading into Mini-PROTEAN® TGX™ Precast Protein Gels, 4–20% (Bio-Rad). Protein was run through the gel at a constant 105 volts, with Precision Plus Protein™ Dual Color Standards, 10–250 kDa (Bio-Rad) included in the first well. Protein was then transferred to a PVDF membrane at 4°C running a constant 100 volts for 1 hour. The PVDF membrane was blocked for 1 hour in TBST (tris-buffered saline with 0.05% Tween) with 3% BSA (bovine serum albumin, Sigma-Aldrich) and then incubated overnight in block solution with rabbit anti-Cdc42bpb (1:200 Novus Biologicals, NBP1-81440) and mouse anti-beta Actin (1:5000 Invitrogen, MA5-15739) at 4°C with agitation. The membrane was then washed with TBST, incubated with HRP-conjugated secondaries, goat anti-rabbit (1:1000 Bio-Techne, HAF008) and goat anti-mouse (1:5000 Invitrogen, G21040), for 1 hour at room temperature and then washed again with TBST. Bands were then visualized using Pierce^TM^ SuperSignal^TM^ West Pico PLUS Chemiluminescent substrate (Thermo Scientific) and imaged with a G:Box Chemi XRQ blot imager (Syngene). Band intensities were then quantified in Fiji (ImageJ) by measuring greyscale intensity, subtracting background (unstained membrane) and normalizing Cdc42bpb band intensity to control β-Actin band intensity. Genotype normalized band intensities were then statistically compared using a one-way ANOVA with multiple comparisons using GraphPad Prism version 10.0.0 for Windows, GraphPad Software, Boston, Massachusetts USA, www.graphpad.com.

### Colorimetric myosin staining

E13.5-E16.5 embryos were collected from *Cdc42bpb^-/+^* timed matings, fixed in 4% PFA overnight at 4°C, then micro-dissected to expose the cranial side of the diaphragm, as previously described (8). Embryos were then bleached for 2 hours at room temperature in Dent’s bleach (54% methanol, 33% H_2_O_2_, 13% DMSO), washed with 100% methanol and stored in Dent’s fix (80% methanol, 20% DMSO) for a minimum of two weeks at 4°C. After extended fixation embryos were washed with PBS, including a one-hour wash at room temperature, then endogenous alkaline phosphatases were inactivated by incubating in 65°C PBS. Embryos were then washed in room temperature PBS, blocked for one hour in 5% goat serum (ThermoFisher Scientific), 20% DMSO and 75% PBS, then transferred to fresh block with Monoclonal Anti-Myosin (skeletal, fast)-Alkaline Phosphatase Sigma #A4335 Mouse IgG1 at a dilution of 1:150 and incubated at room temperature for two days. Embryos were then washed with PBS, then in NTMT (100 mM NaCl, 100 mM Tris-HCl ph 9.5, 50 mM MgCl_2_). Embryos were then stained with a 50% NTMT, 50% 1-Step^TM^ NBT/BCIP substrate solution (Thermo Scientific) for 30 minutes, then post-fixed in 4% PFA/0.1% glutaraldehyde. Stained diaphragms were then imaged on Leica M125 dissecting microscopes and ventral gaps were measured in Fiji (ImageJ), drawing lines between muscle boundaries at the midpoint between the central tendon and mid-ventral body wall. Gap distances were statistically compared across genotypes at each timepoint using one-way ANOVA with multiple comparisons using GraphPad Prism version 10.0.0 for Windows, GraphPad Software, Boston, Massachusetts USA, www.graphpad.com.

### Developing lung immunoffuorescent staining

E13.5 embryos were collected from *Cdc42bpb^-/+^* timed matings, lungs were micro-dissected and fixed for 15 minutes in 4% PFA on ice. Fixed lungs were then washed with PBS and PBS with 0.1% TritonX-100 (PBSTx) (Thermo Scientific), blocked in PBSTx with 10% heat-inactivated sheep serum and incubated with Monoclonal Anti-Cytokeratin, pan antibody produced in mouse (1:250 Sigma-Aldrich, C2931) in block solution overnight at 4°C. Lungs were then washed in PBSTx at 4°C and incubated with anti-MsIgG1 dylight488 secondary antibody at a dilution of 1:250 in block solution in 4°C overnight. Lungs were then washed in PBSTx at 4°C, then imaged on a Leica M165FC fluorescent dissecting microscope. Distal buds were then counted using Fiji (ImageJ), then genotypes were compared statistically using one-way ANOVA with multiple comparison using GraphPad Prism version 10.0.0 for Windows, GraphPad Software, Boston, Massachusetts USA, www.graphpad.com.

### Bulk RNA sequencing and analysis

Embryos were collected from *Cnot1^+/RC23W^* x *Cnot1^+/R623W^* timed matings and diaphragms were micro-dissected as previously described (57), snap frozen in liquid nitrogen and stored at -80°C. Total RNA was isolated from tissue using the Qiagen miRNeasy micro RNA Isolation Kit (Qiagen). Tissues were homogenized in TRIzol (ThermoFisher) using a Pellet Pestle Motor (Kimbal). After the addition of chloroform, the RNA-containing aqueous layer was removed for RNA isolation according to the manufacturer’s protocol. RNA concentration and quality were assessed using the Nanodrop 2000 spectrophotometer (Thermo Scientific) and the RNA 600 Nano Assay (Agilent Technologies). Stranded libraries were constructed using the KAPA RNA Hyper Prep Kit with RiboErase (HMR) (Roche Sequencing and Life Science), according to the manufacturer’s protocol. Briefly, the protocol entails depletion of ribosomal RNA (rRNA), RNA fragmentation, first and second strand cDNA synthesis, ligation of Illumina-specific adapters containing a unique barcode sequence for each library, magnetic bead size selection, and PCR amplification. The quality and concentration of the libraries were assessed using the D5000 ScreenTape (Agilent Technologies) and Qubit dsDNA HS Assay (ThermoFisher), respectively, according to the manufacturers’ instructions. Libraries were sequenced 150 bp paired-end on an Illumina NovaSeq 6000 using the S4 Reagent Kit v1.5. RNA sequencing reads were batch processed using The Jackson Laboratory standard Nextflow RNAseq pipeline (https://github.com/TheJacksonLaboratory/jds-nf-workflows/wiki/RNA-Pipeline-ReadMe), aligning to the mm10 (GRCm38) mouse genome and gencode gene annotation release M25 (GRCm38.6). In brief, sequenced reads were trimmed with Trim Galore (https://github.com/FelixKrueger/TrimGalore), aligned STAR (58) and counts were quantified with Salmon (59) and STRINGTIE (60). Differential gene and transcript analysis was performed with DESeq2 (61), using standard parameters. Statistically significant differentially expressed genes and transcripts per comparison were analyzed with STRING DB (https://string-db.org/) with standard parameters. Alternative polyadenylation between genotypes was determined using APALYZER 3’ UTR analysis standard parameters (42). Volcano plots comparing RED to -Log_10_(P-value) were generated by APALYZER, RED to - Log_10_(Adjusted P-value) were generated using ggplot2 (62). RNA sequencing data is available on GEO at accession number GSE318891.

## Acknowledgments

We thank current and past members of the Murray and Bult groups at The Jackson Laboratory and the Pediatric Surgical Research Laboratory group at Massachusetts General Hospital. We thank Ian Welsh of the Murray group and Julie Wells of the Bult group for editorial input on the manuscript and intellectual input through discussions of data. We thank multiple Scientific Services at The Jackson Laboratory including; Microscopy Services (RRID: SCR_024405), Genetic Engineering Technologies, Genome Technologies Service, Histopathology Sciences Service, and Transgenic Genotyping Services. This work was supported by the NIH Common Fund and the Office of The Director, National Institutes of Health under Award Numbers UM1OD023222 and UM1OD023222-08S3 to S.A.M., R01HD115718 to S.A.M. and F.A.H. E.L.B. was supported by T32HD007065 and F32HD107804. Y.S., P.D., W.K.C., and F.A.H. were supported by P01HD068250. S.P.R. was supported by R03HD112717-01, the Charles H. Hood Childhood Health Award, and the American Lung Association Catalyst Award CA-1052191. Y.L. was supported by Erasmus Traineeship 2024-1-DE01-KA131-HED-000206724. The content is solely the responsibility of the authors and does not necessarily represent the official views of the National Institutes of Health.

## Author Contributions

ELB Conceptualization, Investigation, Formal analysis, Writing-original draft, Funding acquisition

CC Investigation

AW Investigation

KP investigation

AM investigation, formal analysis, Writing – Original Draft

YL investigation

CH investigation

KJS conceptualization, investigation

CB writing-review & editing, funding acquisition

SPR conceptualization, investigation, formal analysis, Writing-Editing

YS formal analysis

PD writing-review & editing, funding acquisition

FAH Conceptualization, Supervision, Investigation, Writing – Editing, Funding acquisition

WKC Writing-review & editing, Project administration

SAM Conceptualization, Supervision, Writing – Original Draft, Funding acquisition

## References

1 Kardon, G., Ackerman, K.G., McCulley, D.J., Shen, Y., Wynn, J., Shang, L., Bogenschutz, E., Sun, X. and Chung, W.K. (2017) Congenital diaphragmatic hernias: from genes to mechanisms to therapies. Dis Model Mech, 10, 955–970.

2 Zani, A., Chung, W.K., Deprest, J., Harting, M.T., Jancelewicz, T., Kunisaki, S.M., Patel, N., Antounians, L., Puligandla, P.S. and Keijzer, R. (2022) Congenital diaphragmatic hernia. Nat Rev Dis Primers, 8, 37.

3 Ackerman, K.G. and Pober, B.R. (2007) Congenital diaphragmatic hernia and pulmonary hypoplasia: new insights from developmental biology and genetics. Am J Med Genet C Semin Med Genet, 145C, 105–108.

4 Longoni, M., High, F.A., Qi, H., Joy, M.P., Hila, R., Coletti, C.M., Wynn, J., Loscertales, M., Shan, L., Bult, C.J., et al. (2017) Genome-wide enrichment of damaging de novo variants in patients with isolated and complex congenital diaphragmatic hernia. Hum Genet, 136, 679–691.

5 Qiao, L., Welch, C.L., Hernan, R., Wynn, J., Krishnan, U.S., Zalieckas, J.M., Buchmiller, T., Khlevner, J., De, A., Farkouh-Karoleski, C., et al. (2024) Common variants increase risk for congenital diaphragmatic hernia within the context of de novo variants. Am J Hum Genet, 111, 2362–2381.

6 Bogenschutz, E.L., Fox, Z.D., Farrell, A., Wynn, J., Moore, B., Yu, L., Aspelund, G., Marth, G., Yandell, M., Shen, Y., et al. (2020) Deep whole-genome sequencing of multiple proband tissues and parental blood reveals the complex genetic etiology of congenital diaphragmatic hernias. HGG Adv, 1.

7 Perry, S.F., Similowski, T., Klein, W. and Codd, J.R. (2010) The evolutionary origin of the mammalian diaphragm. Respir Physiol Neurobiol, 171, 1–16.

8 Merrell, A.J., Ellis, B.J., Fox, Z.D., Lawson, J.A., Weiss, J.A. and Kardon, G. (2015) Muscle connective tissue controls development of the diaphragm and is a source of congenital diaphragmatic hernias. Nat Genet, 47, 496–504.

9 Sefton, E.M., Gallardo, M. and Kardon, G. (2018) Developmental origin and morphogenesis of the diaphragm, an essential mammalian muscle. Dev Biol, 440, 64–73.

10 Allan, D.W. and Greer, J.J. (1997) Embryogenesis of the phrenic nerve and diaphragm in the fetal rat. J Comp Neurol, 382, 459–468.

11 Babiuk, R.P. and Greer, J.J. (2002) Diaphragm defects occur in a CDH hernia model independently of myogenesis and lung formation. Am J Physiol Lung Cell Mol Physiol, 283, L1310–1314.

12 Paris, N.D., Coles, G.L. and Ackerman, K.G. (2015) Wt1 and beta-catenin cooperatively regulate diaphragm development in the mouse. Dev Biol, 407, 40–56.

13 Carmona, R., Canete, A., Cano, E., Ariza, L., Rojas, A. and Munoz-Chapuli, R. (2016) Conditional deletion of WT1 in the septum transversum mesenchyme causes congenital diaphragmatic hernia in mice. Elife, 5.

14 Sefton, E.M., Gallardo, M., Tobin, C.E., Collins, B.C., Colasanto, M.P., Merrell, A.J. and Kardon, G. (2022) Fibroblast-derived Hgf controls recruitment and expansion of muscle during morphogenesis of the mammalian diaphragm. Elife, 11.

15 Guevara, G., Wild, K.T., Keijzer, R. and McCulley, D.J. (2025) Developmental pathophysiology and genetic contributions in CDH. Semin Fetal Neonatal Med, 30, 101652.

16 Unbekandt, M. and Olson, M.F. (2014) The actin-myosin regulatory MRCK kinases: regulation, biological functions and associations with human cancer. J Mol Med (Berl), 92, 217–225.

17 Heikkila, T., Wheatley, E., Crighton, D., Schroder, E., Boakes, A., Kaye, S.J., Mezna, M., Pang, L., Rushbrooke, M., Turnbull, A. et al. (2011) Co-crystal structures of inhibitors with MRCKbeta, a key regulator of tumor cell invasion. PLoS One, 6, e24825.

18 Pichaud, F., Walther, R.F. and Nunes de Almeida, F. (2019) Regulation of Cdc42 and its effectors in epithelial morphogenesis. J Cell Sci, 132.

19 Vicente-Manzanares, M., Ma, X., Adelstein, R.S. and Horwitz, A.R. (2009) Non-muscle myosin II takes centre stage in cell adhesion and migration. Nat Rev Mol Cell Biol, 10, 778–790.

20 Huo, L., Wen, W., Wang, R., Kam, C., Xia, J., Feng, W. and Zhang, M. (2011) Cdc42-dependent formation of the ZO-1/MRCKbeta complex at the leading edge controls cell migration. EMBO J, 30, 665–678.

21 Chilton, I., Okur, V., Vitiello, G., Selicorni, A., Mariani, M., Goldenberg, A., Husson, T., Campion, D., Lichtenbelt, K.D., van Gassen, K., et al. (2020) De novo heterozygous missense and loss-of-function variants in CDC42BPB are associated with a neurodevelopmental phenotype. Am J Med Genet A, 182, 962–973.

22 Bartlam, M. and Yamamoto, T. (2010) The structural basis for deadenylation by the CCR4-NOT complex. Protein Cell, 1, 443–452.

23 Chalabi Hagkarim, N. and Grand, R.J. (2020) The Regulatory Properties of the Ccr4-Not Complex. Cells, 9.

24 Ito, K., Takahashi, A., Morita, M., Suzuki, T. and Yamamoto, T. (2011) The role of the CNOT1 subunit of the CCR4-NOT complex in mRNA deadenylation and cell viability. Protein Cell, 2, 755–763.

25 Yamaguchi, T., Suzuki, T., Sato, T., Takahashi, A., Watanabe, H., Kadowaki, A., Natsui, M., Inagaki, H., Arakawa, S., Nakaoka, S. et al. (2018) The CCR4-NOT deadenylase complex controls Atg7-dependent cell death and heart function. Sci Signal, 11.

26 Takahashi, A., Suzuki, T., Soeda, S., Takaoka, S., Kobori, S., Yamaguchi, T., Mohamed, H.M.A., Yanagiya, A., Abe, T., Shigeta, M. et al. (2020) The CCR4-NOT complex maintains liver homeostasis through mRNA deadenylation. Life Sci Alliance, 3.

27 Kruszka, P., Berger, S.I., Weiss, K., Everson, J.L., Martinez, A.F., Hong, S., Anyane-Yeboa, K., Lipinski, R.J. and Muenke, M. (2019) A CCR4-NOT Transcription Complex, Subunit 1, CNOT1, Variant Associated with Holoprosencephaly. Am J Hum Genet, 104, 990–993.

28 Vissers, L., Kalvakuri, S., de Boer, E., Geuer, S., Oud, M., van Outersterp, I., Kwint, M., Witmond, M., Kersten, S., Polla, D.L., et al. (2020) De Novo Variants in CNOT1, a Central Component of the CCR4-NOT Complex Involved in Gene Expression and RNA and Protein Stability, Cause Neurodevelopmental Delay. Am J Hum Genet, 107, 164–172.

29 Dong, Y., Li, W., Meng, J., Wang, P., Sun, M., Zhou, F., Li, D., Shu, J. and Cai, C. (2023) Pathogenicity analysis and splicing rescue of a classical splice site variant (c.1343+1G>T) of CNOT1 gene associated with neurodevelopmental disorders. Am J Med Genet A, 191, 2775–2782.

30 De Franco, E., Watson, R.A., Weninger, W.J., Wong, C.C., Flanagan, S.E., Caswell, R., Green, A., Tudor, C., Lelliott, C.J., Geyer, S.H., et al. (2019) A Specific CNOT1 Mutation Results in a Novel Syndrome of Pancreatic Agenesis and Holoprosencephaly through Impaired Pancreatic and Neurological Development. Am J Hum Genet, 104, 985–989.

31 Pfeufer, A., Sanna, S., Arking, D.E., Muller, M., Gateva, V., Fuchsberger, C., Ehret, G.B., Orru, M., Pattaro, C., Kottgen, A. et al. (2009) Common variants at ten loci modulate the QT interval duration in the QTSCD Study. Nat Genet, 41, 407–414.

32 Newton-Cheh, C., Eijgelsheim, M., Rice, K.M., de Bakker, P.I., Yin, X., Estrada, K., Bis, J.C., Marciante, K., Rivadeneira, F., Noseworthy, P.A., et al. (2009) Common variants at ten loci influence QT interval duration in the QTGEN Study. Nat Genet, 41, 399–406.

33 Gutierrez-Camino, A., Lopez-Lopez, E., Martin-Guerrero, I., Pinan, M.A., Garcia-Miguel, P., Sanchez-Toledo, J., Carbone Baneres, A., Uriz, J., Navajas, A. and Garcia-Orad, A. (2014) Noncoding RNA-related polymorphisms in pediatric acute lymphoblastic leukemia susceptibility. Pediatr Res, 75, 767–773.

34 Qiao, L., Xu, L., Yu, L., Wynn, J., Hernan, R., Zhou, X., Farkouh-Karoleski, C., Krishnan, U.S., Khlevner, J., De, A., et al. (2021) Rare and de novo variants in 827 congenital diaphragmatic hernia probands implicate LONP1 as candidate risk gene. Am J Hum Genet, 108, 1964–1980.

35 Dickinson, M.E., Flenniken, A.M., Ji, X., Teboul, L., Wong, M.D., White, J.K., Meehan, T.F., Weninger, W.J., Westerberg, H., Adissu, H. et al. (2016) High-throughput discovery of novel developmental phenotypes. Nature, 537, 508–514.

36 Yushkevich, P.A., Piven, J., Hazlett, H.C., Smith, R.G., Ho, S., Gee, J.C. and Gerig, G. (2006) User-guided 3D active contour segmentation of anatomical structures: significantly improved efficiency and reliability. Neuroimage, 31, 1116–1128.

37 Guimier, A., Gabriel, G.C., Bajolle, F., Tsang, M., Liu, H., Noll, A., Schwartz, M., El Malti, R., Smith, L.D., Klena, N.T. et al. (2015) MMP21 is mutated in human heterotaxy and is required for normal left-right asymmetry in vertebrates. Nat Genet, 47, 1260–1263.

38 Liu, X., Yagi, H., Saeed, S., Bais, A.S., Gabriel, G.C., Chen, Z., Peterson, K.A., Li, Y., Schwartz, M.C., Reynolds, W.T. et al. (2017) The complex genetics of hypoplastic left heart syndrome. Nat Genet, 49, 1152–1159.

39 Szklarczyk, D., Kirsch, R., Koutrouli, M., Nastou, K., Mehryary, F., Hachilif, R., Gable, A.L., Fang, T., Doncheva, N.T., Pyysalo, S. et al. (2023) The STRING database in 2023: protein-protein association networks and functional enrichment analyses for any sequenced genome of interest. Nucleic Acids Res, 51, D638–D646.

40 Smith, C.L. and Eppig, J.T. (2009) The mammalian phenotype ontology: enabling robust annotation and comparative analysis. Wiley Interdiscip Rev Syst Biol Med, 1, 390–399.

41 Baldarelli, R.M., Smith, C.L., Ringwald, M., Richardson, J.E., Bult, C.J. and Mouse Genome Informatics, G. (2024) Mouse Genome Informatics: an integrated knowledgebase system for the laboratory mouse. Genetics, 227.

42 Wang, R. and Tian, B. (2020) APAlyzer: a bioinformatics package for analysis of alternative polyadenylation isoforms. Bioinformatics, 36, 3907–3909.

43 Robinson, J.T., Thorvaldsdottir, H., Winckler, W., Guttman, M., Lander, E.S., Getz, G. and Mesirov, J.P. (2011) Integrative genomics viewer. Nat Biotechnol, 29, 24–26.

44 Petit, F., Longoni, M., Wells, J., Maser, R.S., Bogenschutz, E.L., Dysart, M.J., Contreras, H.T.M., Frenois, F., Pober, B.R., Clark, R.D. et al. (2023) PLS3 missense variants affecting the actin-binding domains cause X-linked congenital diaphragmatic hernia and body-wall defects. Am J Hum Genet, 110, 1787–1803.

45 UniProt, C. (2025) UniProt: the Universal Protein Knowledgebase in 2025. Nucleic Acids Res, 53, D609–D617.

46 Hagkarim, N.C., Hajkarim, M.C., Suzuki, T., Fujiwara, T., Winkler, G.S., Stewart, G.S. and Grand, R.J. (2023) Disruption of the Mammalian Ccr4-Not Complex Contributes to Transcription-Mediated Genome Instability. Cells, 12.

47 Stowell, J.A.W., Yu, C.W.H., Chen, Z.A., DeBell, L.K., Lee, G., Morgan, T., Sinn, L., Agnello, S., O’Reilly, F.J., Rappsilber, J., et al. (2025) Phosphorylation-dependent tuning of mRNA deadenylation rates. Nat Struct Mol Biol, in press.

48 Tuzovic, L., Yu, L., Zeng, W., Li, X., Lu, H., Lu, H.M., Gonzalez, K.D. and Chung, W.K. (2013) A human de novo mutation in MYH10 phenocopies the loss of function mutation in mice. Rare Dis, 1, e26144.

49 Tevosian, S.G., Deconinck, A.E., Tanaka, M., Schinke, M., Litovsky, S.H., Izumo, S., Fujiwara, Y. and Orkin, S.H. (2000) FOG-2, a cofactor for GATA transcription factors, is essential for heart morphogenesis and development of coronary vessels from epicardium. Cell, 101, 729–739.

50 Kuo, C.T., Morrisey, E.E., Anandappa, R., Sigrist, K., Lu, M.M., Parmacek, M.S., Soudais, C. and Leiden, J.M. (1997) GATA4 transcription factor is required for ventral morphogenesis and heart tube formation. Genes Dev, 11, 1048–1060.

51 Molkentin, J.D., Lin, Q., Duncan, S.A. and Olson, E.N. (1997) Requirement of the transcription factor GATA4 for heart tube formation and ventral morphogenesis. Genes Dev, 11, 1061–1072.

52 Garcia Rivas, J.F., Applin, N.H.M., Albrechtsen, J.F.P., Ghazanfari, A., Doschak, M. and Clugston, R.D. (2025) Mesenchymal retinoic acid signaling is required for normal diaphragm development in mice. FASEB J, 39, e70381.

53 Kaplan, M.M., Zeidler, M., Knapp, A., Holzl, M., Kress, M., Fritsch, H., Krogsdam, A. and Flucher, B.E. (2024) Spatial transcriptomics in embryonic mouse diaphragm muscle reveals regional gradients and subdomains of developmental gene expression. iScience, 27, 110018.

54 Hsu, P.D., Scott, D.A., Weinstein, J.A., Ran, F.A., Konermann, S., Agarwala, V., Li, Y., Fine, E.J., Wu, X., Shalem, O. et al. (2013) DNA targeting specificity of RNA-guided Cas9 nucleases. Nat Biotechnol, 31, 827–832.

55 Doench, J.G., Fusi, N., Sullender, M., Hegde, M., Vaimberg, E.W., Donovan, K.F., Smith, I., Tothova, Z., Wilen, C., Orchard, R. et al. (2016) Optimized sgRNA design to maximize activity and minimize off-target effects of CRISPR-Cas9. Nat Biotechnol, 34, 184–191.

56 Modzelewski, A.J., Chen, S., Willis, B.J., Lloyd, K.C.K., Wood, J.A. and He, L. (2018) Efficient mouse genome engineering by CRISPR-EZ technology. Nat Protoc, 13, 1253–1274.

57 Bogenschutz, E.L., Sefton, E.M. and Kardon, G. (2020) Cell culture system to assay candidate genes and molecular pathways implicated in congenital diaphragmatic hernias. Dev Biol, 467, 30–38.

58 Dobin, A., Davis, C.A., Schlesinger, F., Drenkow, J., Zaleski, C., Jha, S., Batut, P., Chaisson, M. and Gingeras, T.R. (2013) STAR: ultrafast universal RNA-seq aligner. Bioinformatics, 29, 15–21.

59 Patro, R., Duggal, G., Love, M.I., Irizarry, R.A. and Kingsford, C. (2017) Salmon provides fast and bias-aware quantification of transcript expression. Nat Methods, 14, 417–419.

60 Pertea, M., Pertea, G.M., Antonescu, C.M., Chang, T.C., Mendell, J.T. and Salzberg, S.L. (2015) StringTie enables improved reconstruction of a transcriptome from RNA-seq reads. Nat Biotechnol, 33, 290–295.

61 Love, M.I., Huber, W. and Anders, S. (2014) Moderated estimation of fold change and dispersion for RNA-seq data with DESeq2. Genome Biol, 15, 550.

62 Wickham, H. and Sievert, C. (2016) ggplot2 : Elegant Graphics for Data Analysis. Springer International Publishing, Cham.

