## Supplementary Figures for "Modeling patient variants of *Cnot1* and *Cdc42bpb* results in distinct forms of congenital diaphragmatic hernia in mice"

**Supplementary information**

This file includes:

Supplementary Figures 1-6

Descriptions of Supplementary Tables 1-7

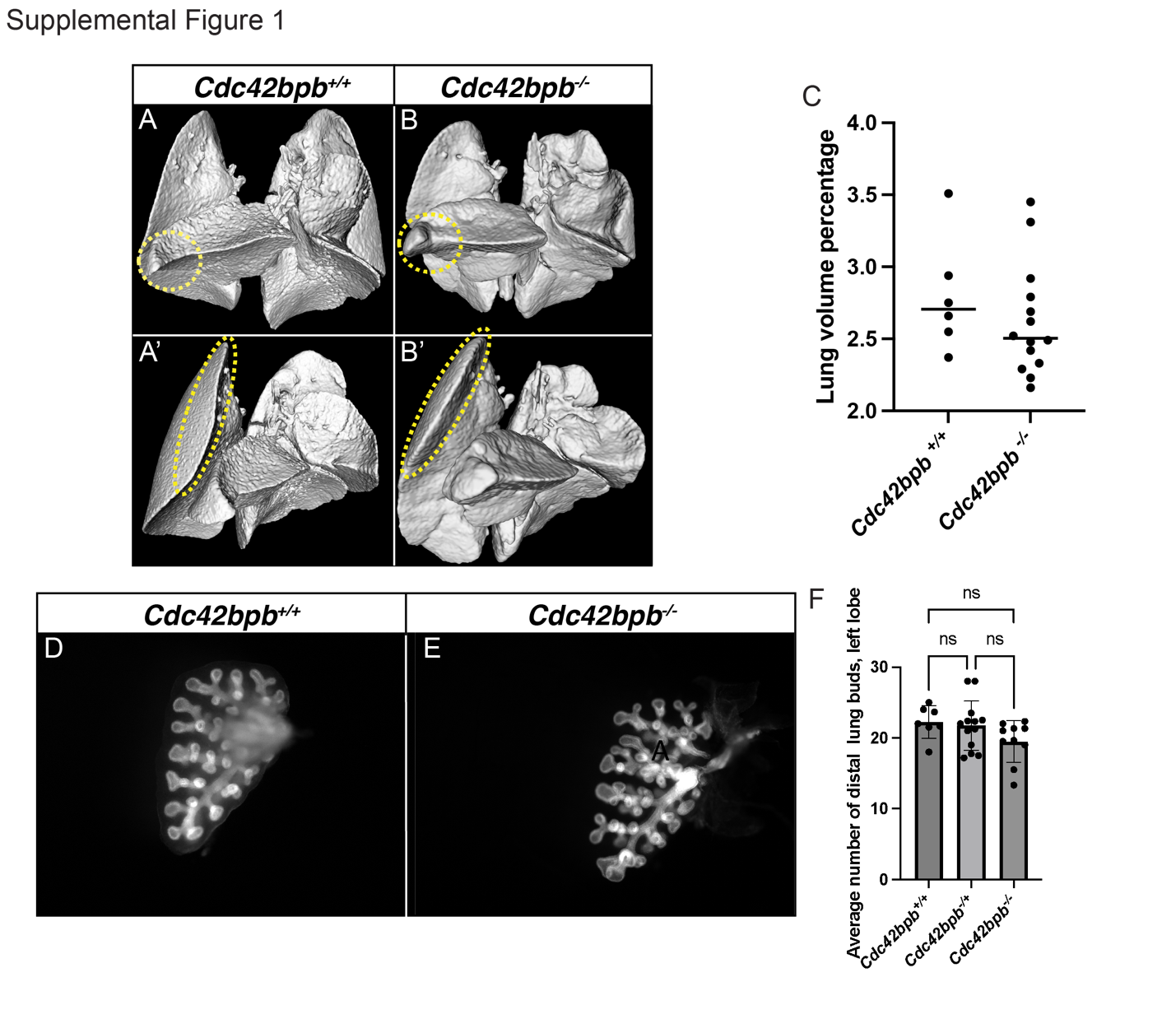

**Supplemental Figure 1**: Lung morphology of *Cdc42bpb^-/-^* mutants. Representative μCT surface renderings of E18.5 lungs A) *Cdc42bpb^+/+^* and B) *Cdc42bpb^-/-^* mutants, showing dysmorphic features including blunted distal edges of lung lobes and accessory lobe horn (yellow dashed-lines). C) Comparison of lung volume percentage (rendered lung volume/whole embryo render volume) between *Cdc42bpb^+/+^* and *Cdc42bpb^-/-^*, which were not significant (Student’s t-test). D, E) Whole mount pan-cytokeratin immunofluorescence of E13.5 left lung lobes, show similar lung branching pattern between *Cdc42bpb^+/+^* controls and *Cdc42bpb^-/-^* mutants. F) Quantitation of number of distal buds for each genotype (not significant by one-way ANOVA).

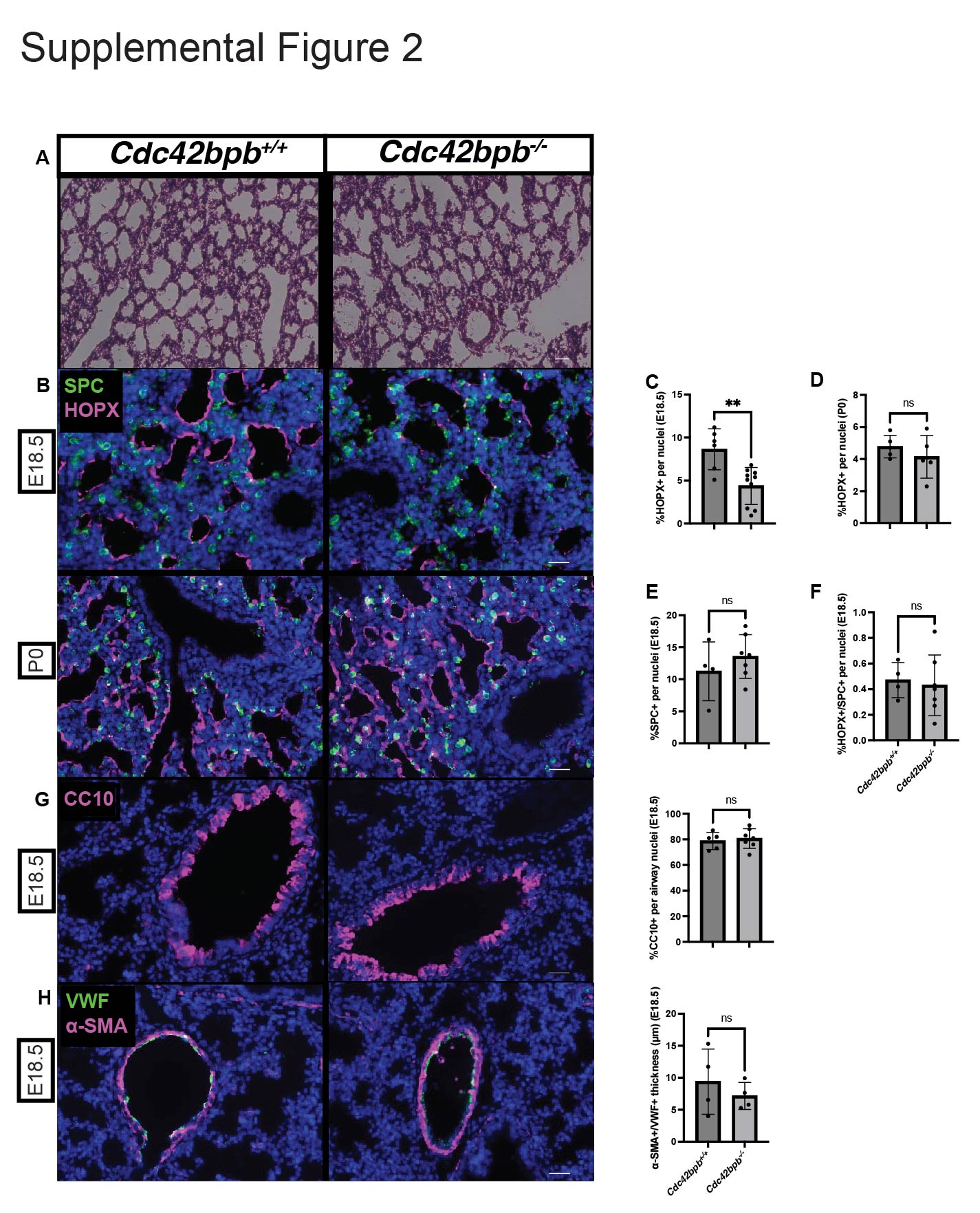

**Supplemental Figure 2**: E18.5 *Cdc42bpb^-/-^* mice exhibit a delay in ATI cell differentiation. A) H&E stainings of E18.5 *Cdc42bpb^+/+^* and *Cdc42bpb^-/-^* mice. B) Immunostainings of alveolar subpopulations in *Cdc42bpb^+/+^* and *Cdc42bpb^-/-^* mice at E18.5 and P0. C) Quantification of E18.5 HOPX+ ATI cell population reduction in *Cdc42bpb^+/+^* and *Cdc42bpb^-/-^* mice (p= 0.0051, T-test). D) Quantification of P0 HOPX+ ATI population in *Cdc42bpb^+/+^* and *Cdc42bpb^-/-^* mice (p=0.3784). E) Quantification of E18.5 SPC staining of the ATII population in *Cdc42bpb^+/+^* and *Cdc42bpb^-/-^* mice (p=0.4238, T-test). F) Quantification of E18.5 double-positive ATI/ATII transitional cell population in *Cdc42bpb^+/+^* and *Cdc42bpb^-/-^* mice (p=0.7310). G) Immunostainings and quantification of CC10+ club cell populations of E18.5 *Cdc42bpb^+/+^* and *Cdc42bpb^-/-^* mice (p=0.6557). H) Vascular smooth muscle and endothelial cell immunostainings with quantification of blood vessel thickness in E18.5 *Cdc42bpb^+/+^* and *Cdc42bpb^-/-^* mice (p=0.4617). **=p<0.01 by Student’s t-test. Scale bar – 25μm

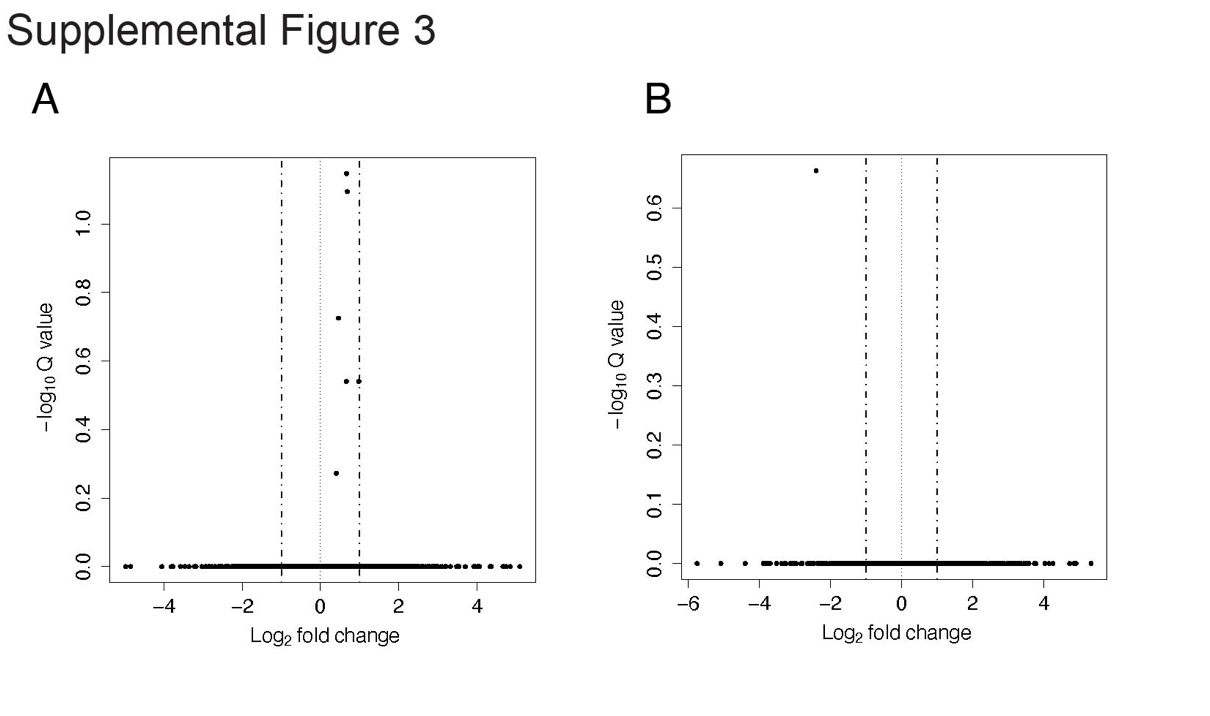

**Supplemental Figure 3**: No differentially expressed genes are found between *Cnot1* mutant embryos and *Cnot1^+/+^*controls at E13.5. Volcano plots of differentially expressed genes between *Cnot1^+/+^* and A) *Cnot1^+/R623W^* or B) *Cnot1^R623W/R623W^* diaphragms. No genes pass p-adjusted (Q) value threshold of 0.05 in either comparison.

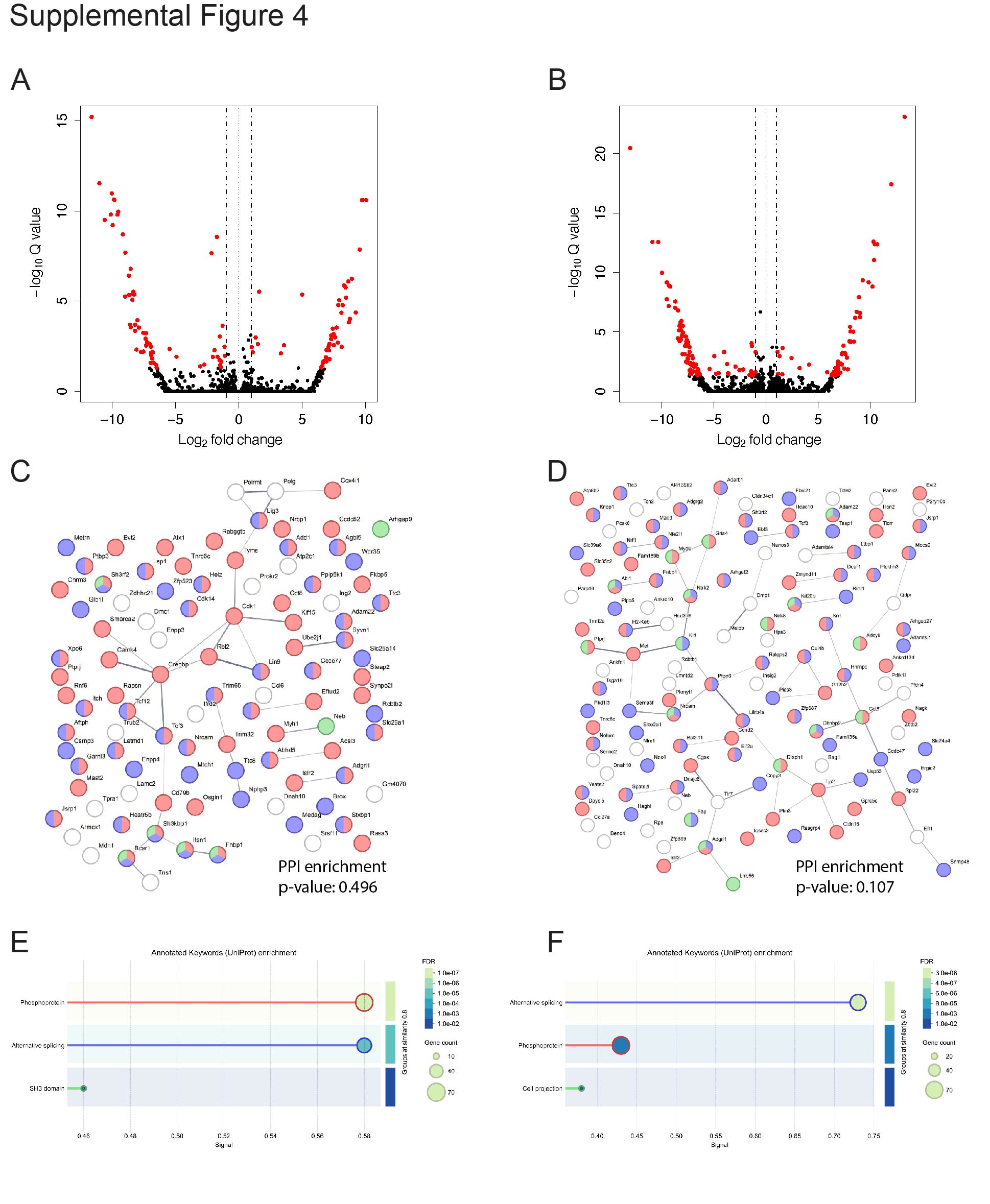

**Supplemental Figure 4**: Transcript differential expression analysis of E13.5 diaphragms. Volcano plots of transcript differential analysis between A) *Cnot1^+/+^* vs. *Cnot1^+/R623W^* and B) *Cnot1^+/+^* vs. *Cnot1^R623W/R623W^*. Transcripts (datapoints) with absolute log_2_ fold change greater than 1 and p-adjusted (Q) values less than 0.05 highlighted in red. STRING gene networks of significant differentially expressed transcripts from C) *Cnot1^+/+^* vs. *Cnot1^+/R623W^* and D) *Cnot1^+/+^* vs. *Cnot1^R623W/R623W^* comparisons. Lines between nodes denote interaction between genes with STRING interaction confidence scores greater than or equal to 0.4. E) and F) show Uniprot terms enriched in transcript networks for C) and D), respectively, by STRING GO analysis. Genes belonging to each Uniprot term are shown fully or partially colored in C) and D) as terms in E) and F).

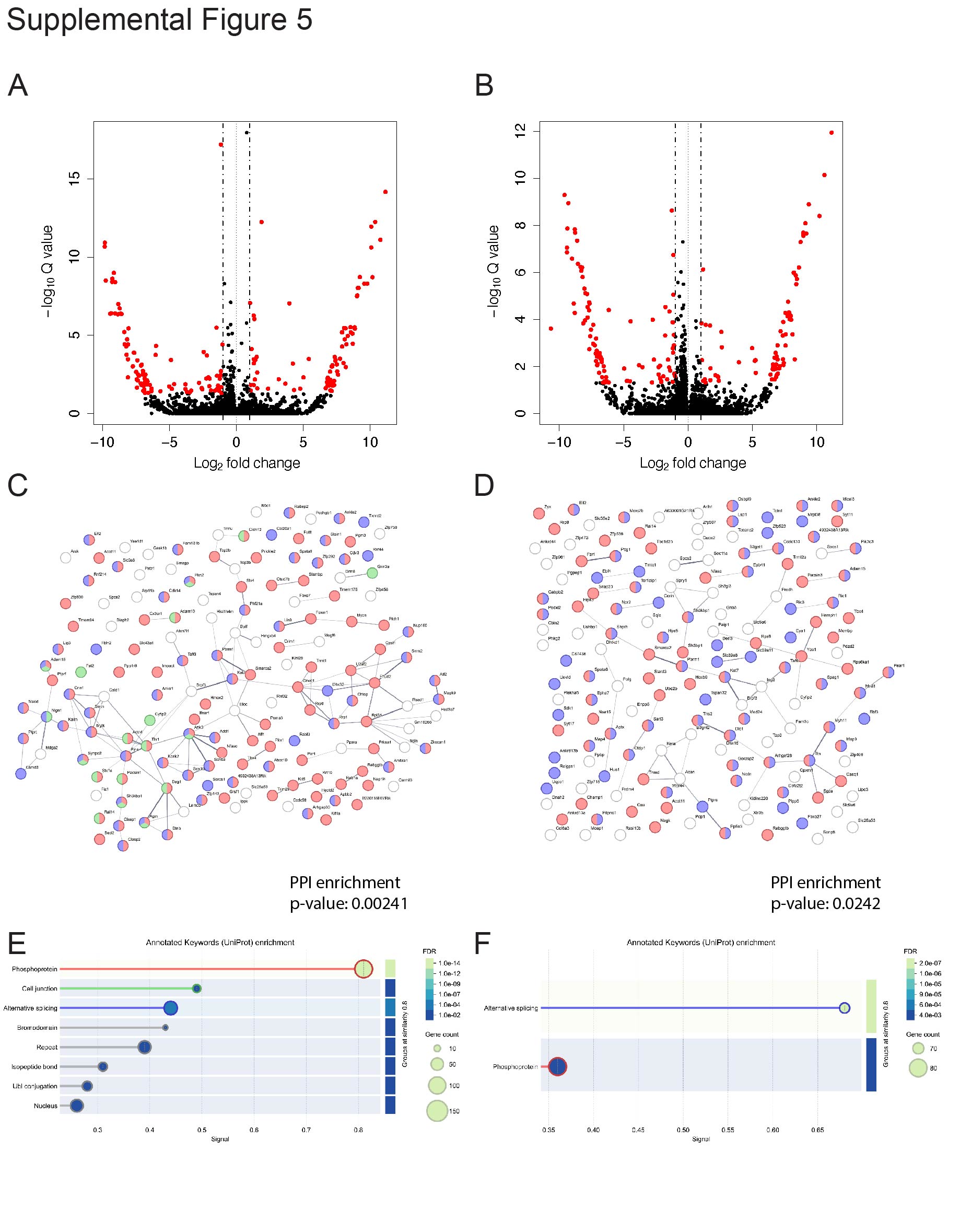

**Supplemental Figure 5**: Transcript differential expression analysis of E14.5 diaphragms. Volcano plots of transcript differential analysis between A) *Cnot1^+/+^* vs. *Cnot1^+/R623W^* and B) *Cnot1^+/+^* vs. *Cnot1^R623W/R623W^*. Transcripts (datapoints) with absolute log_2_ fold change greater than 1 and p-adjusted (Q) values less than 0.05 highlighted in red. STRING gene networks of significant differentially expressed transcripts from C) *Cnot1^+/+^* vs. *Cnot1^+/R623W^* and D) *Cnot1^+/+^* vs. *Cnot1^R623W/R623W^* comparisons. Lines between nodes denote interaction between genes with STRING interaction confidence scores greater than or equal to 0.4. E) and F) show Uniprot terms enriched in transcript networks for C) and D), respectively, by STRING GO analysis. Genes belonging to each Uniprot term are shown fully or partially colored in C) and D) as terms in E) and F).

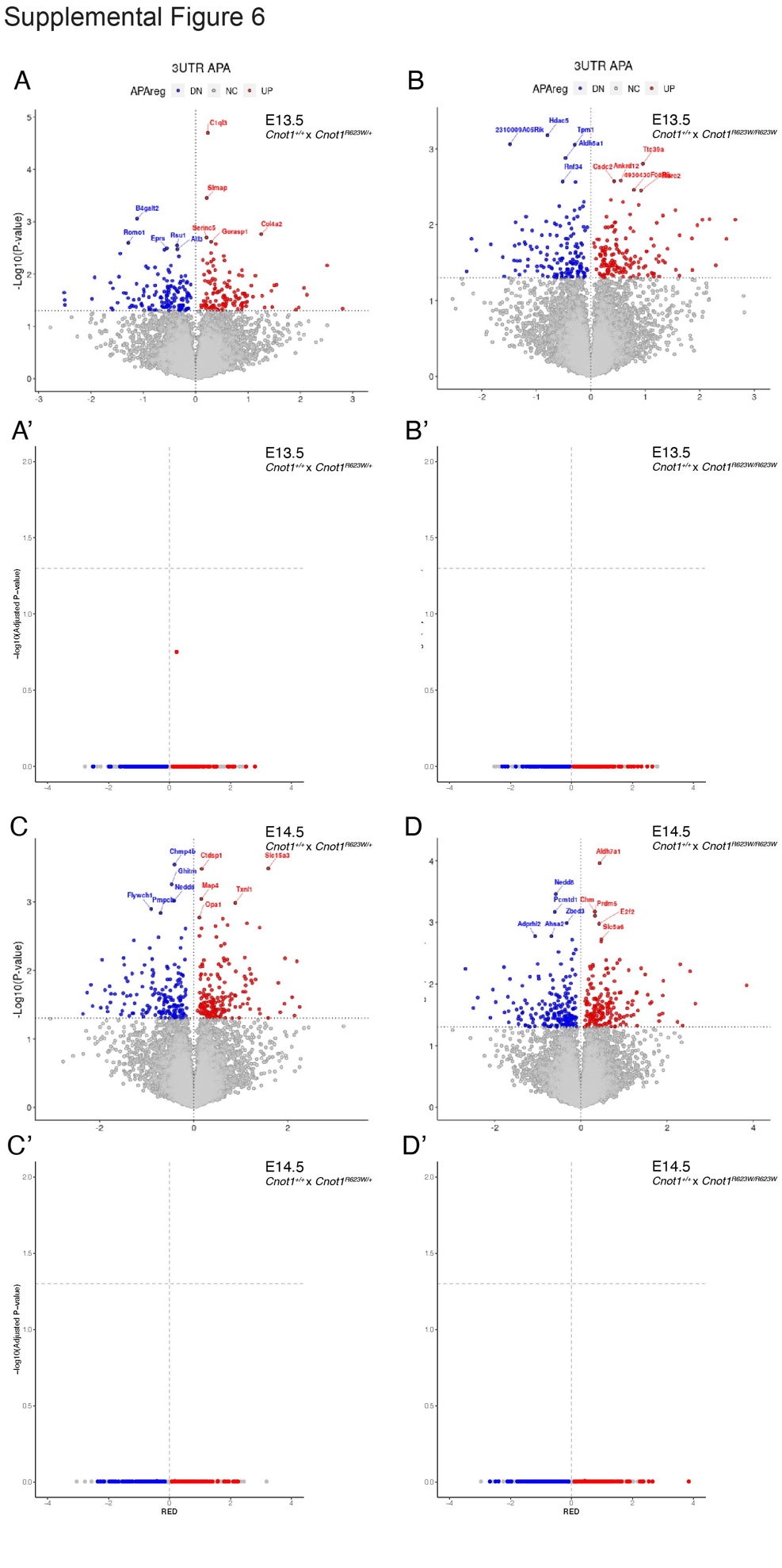

**Supplemental Figure 6**: Alternative polyadenylation analysis. Genes with differential polyadenylation comparing mutant *Cnot1* samples against wildtype at E13.5 (A-B) and E14.5 (C-D). Apalyzer generated APA relative expression score differences (RED) between genotypes compared are plotted on the X-axis (A-D). Normalized -log_10_ of either Apalyzer generated P-value (A, B, C ,D) or multiple test corrected p-adjusted value (A’, B’, C’, D’), as determined by Student’s T-test with Bonferroni correction, plotted on Y-axis. Red datapoints represent genes with increased 3’ UTR polyadenylation (UP), blue datapoints with decreased 3’ UTR polyadenylation (DN) and grey datapoints represent non-significantly changed based on p-value <0.05 (NC). Top ten differential genes labeled in A, B, C, and D.

**Supplemental Table 1**: ICE generated CRISPR editing scores for C>T p.T1179M *Cdc42bpb* F0 embryos.

**Supplemental Table 2**: DESeq2 results of gene level comparisons between RNA-sequencing samples.

**Supplemental Table 3**: STRING DB significant ontology terms for each set of significantly differentially expressed genes across sample comparisons.

**Supplemental Table 4**: DESeq2 results of transcript level comparisons between RNA-sequencing samples.

**Supplemental Table 5**: STRING DB significant ontology terms for each set of significantly differentially expressed transcripts across sample comparisons.

**Supplemental Table 6**: Apalzyer results comparing *Cnot1^+/+^* to either *Cnot1^R623W/+^*or *Cnot^R623W/R623W^* at either E13.5 or E14.5.

**Supplemental Table 7**: List of oligonucleotide sequences used in this study.
